# Super-resolved spatial organization of the nucleolar transcriptome

**DOI:** 10.64898/2026.05.19.726041

**Authors:** Xinqi Fan, Chien-Chien Tan, Joshua H. Mu, Sheng-Min Chiang, Yu Xiao, Chuan He, Isaac T.S. Li, Wei-Sheng Wu, Jingyi Fei

**Affiliations:** Department of Biochemistry and Molecular Biology, The University of Chicago, Chicago, IL 60637, USA; Department of Electrical Engineering, National Cheng Kung University, Tainan 701, Taiwan; Illinois Mathematics and Science Academy, Aurora, IL 60506, USA; Department of Chemistry, The University of Chicago, Chicago, IL 60637, USA; Institute for Biophysical Dynamics, The University of Chicago, Chicago, IL 60637, USA; Howard Hughes Medical Institute, Chicago, IL 60637, USA; Department of Chemistry, University of British Columbia, Kelowna, BC V1V 1V7, Canada

## Abstract

Membraneless organelles (MLOs) often exhibit internal architecture, yet whether the local transcriptome differentially partitions across MLO subdomains remains largely uncharacterized. Here we combine super-resolution imaging with in situ reverse transcription-based sequencing to profile transcriptomes within MLO subdomains. Using the human tripartite nucleolus as a model system, we identify distinct RNA populations in the fibrillar center (FC), dense fibrillar component (DFC), and granular component (GC). Pre-rRNA processing intermediates demonstrate a layered progression across nucleolar subdomains, reflecting the temporal order of the processing steps. Processing steps involved in large-small subunit separation show increased retention in the DFC in highly differentiated cells. Mature small nucleolar RNAs (snoRNAs) are preferentially enriched in the DFC and spatially segregated from their precursor transcripts. Many non-snoRNA-related transcripts, often derived from nucleolus-proximal genes, show modest enrichment in the GC. These results illustrate functional RNA organization across nucleolar subdomains and provide a framework for nanoscale transcriptome mapping of biomolecular condensates.

---

Membraneless organelles (MLOs) are prevalent in eukaryotic cells and widely involved in vital biological activities ^1,2^. Many MLOs contain internal subdomains, often characterized by enrichment of specific protein or RNA components ^2^. A prominent example is the nucleolus, which displays a tripartite structure in mammalian cells, with three layers of fibrillar center (FC), dense fibrillar component (DFC), and granular component (GC) from the core to the periphery (Fig. 1a) ^3–5^. Such internal organization is functionally linked to different stages of ribosome biogenesis. Specifically, FC contains rDNA genes, where active transcription of rDNA occurs at the interface of FC and DFC. Co-transcriptional pre-rRNA processing and modification take place in DFC, and late processing steps and assembly with ribosomal proteins take place in GC ^3,4,6^. In nuclear speckles, the scaffold proteins SON and SRRM2 form the core region, and certain long noncoding RNA (lncRNA) *MALAT1* and other spliceosomal components are distributed at the outer shell ^7^. Similar core-shell organizations have been observed in paraspeckles ^8^, stress granules ^9^, and PML bodies ^10^. Despite the prevalence of internal organizations within MLOs, their functional importance is largely unclear.

**Fig. 1.**
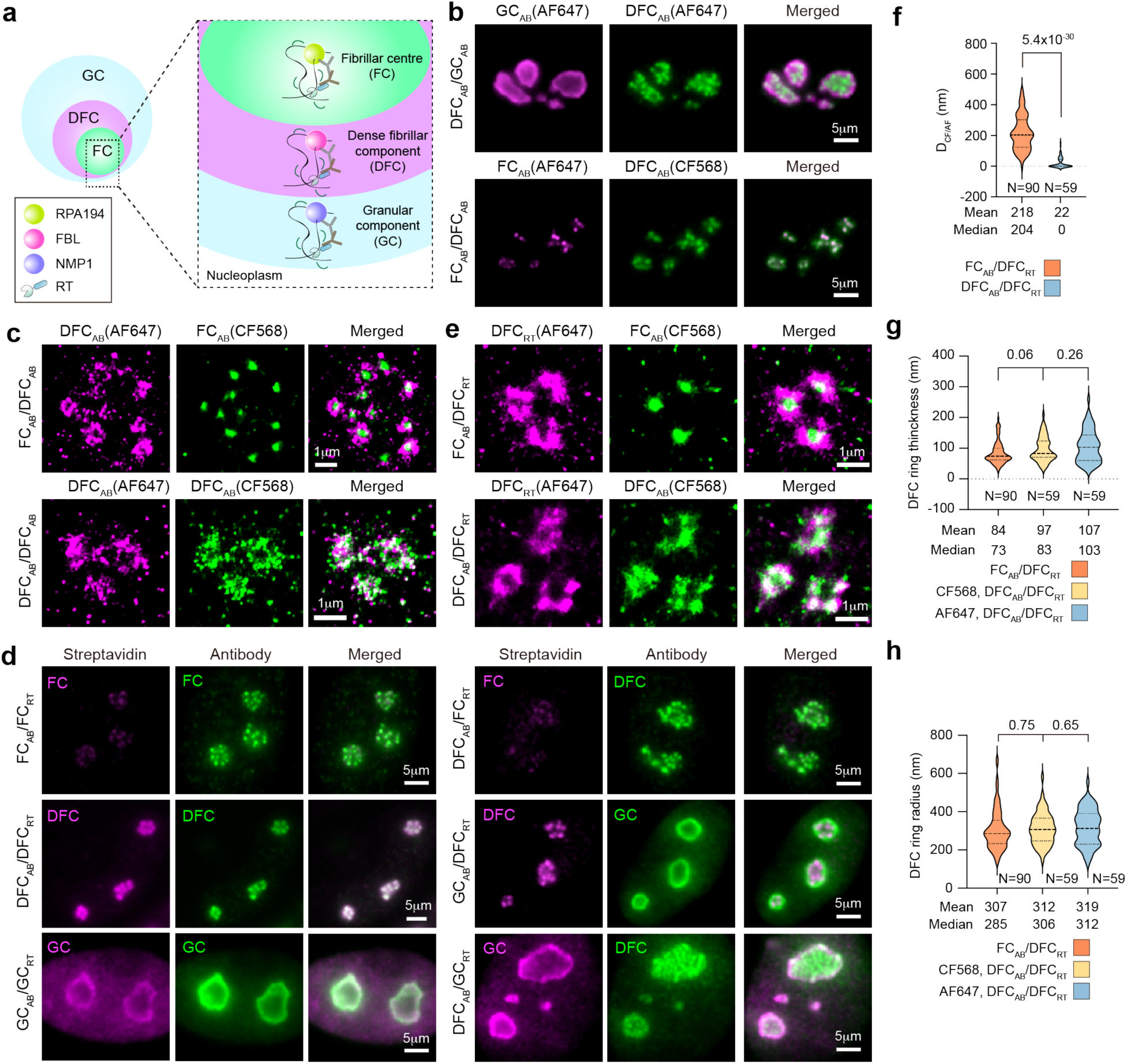
Transcriptome profiling of nucleolar subdomains. **a**, Scheme of in situ reverse transcription used for transcriptome profiling in nucleolar subdomains. **b**, Representative epi-fluorescence images of nucleoli with each domain immunostained in HFF cells. **c**, Representative SMLM images of DF/DFC units by immunostaining within one nucleolus in HFF cells. **d**, Representative epi-fluorescence images of in situ RT-generated biotin-cDNAs stained by AF647-labeled streptavidin, together with each marker protein stained by CF568-labeled antibody in different nucleolar subdomains. Biotin signals in both FC and DFC regions were acquired under identical imaging conditions and displayed with the same contrast settings. Biotin signals in GC were imaged using a lower laser power to avoid image oversaturation. **e**, Representative SMLM images of biotin-cDNA and marker protein in FC/DFC units. Biotin-cDNAs were generated in DFC, while marker proteins of FC or DFC were simultaneously stained with the corresponding antibodies. **f**, Violin plot for cross-correlation distance between AF647 and CF568 channels. **g**, Violin plot of thickness of DFC ring, estimated by autocorrelation distance, *D*_AF/AF, 1_ or *D*_CF/CF, 1_ (Supplementary Text). **h,** Violin plot of DFC radius, estimated by autocorrelation distance, *D*_AF/AF, 2_ or *D*_CF/CF, 2_. The fitted correlation distance (in the unit of pixels) was multiplied by the effective pixel size (26 nm). N in (f-h) reports the number of FC/DFC units in each plot, collected from 7 cells from two biological replicates. *P*-values were all calculated by unpaired two-tailed t test.

Dynamic localization of RNAs to these MLOs is often critically linked to their biological functions. However, prior transcriptomic analyses have largely treated each MLO as a single entity ^11–13^, without resolving subdomain-level organization. Whether the local transcriptome partitions across subdomains of an MLO remains unknown. Approaches that integrate transcriptome profiling with subdomain-level spatial resolution are needed to directly test how intra-MLO RNA localization contributes to MLO function.

We reasoned that a previously developed in situ reverse transcription (RT)-based sequencing method, ARTR-seq, can provide sufficient spatial resolution for subdomain transcriptome profiling in MLOs. In ARTR-seq, a fusion protein (pAG-RTase), which consists of a reverse transcriptase (RTase) and a protein A/G (pAG), is generated, ^14^. After immunostaining of the marker proteins of an MLO of interest, the pAG-RTase can bind to both the primary and secondary antibodies to perform reverse transcription in situ ^15^. The theoretical maximum distance between the targeted protein and the RTase is estimated to be 50 nm, which matches the dimension of subdomains of known MLOs.

Here, we combine super-resolution imaging with ARTR-seq to profile transcriptomes within MLO subdomains. Using the human tripartite nucleolus as a model system, we identify distinct RNA populations in the FC, DFC, and GC subdomains and uncover a layered progression of pre-rRNA processing across nucleolar subdomains. Our approach establishes a framework for nanoscale transcriptome mapping of biomolecular condensates.

## Results

### ARTR-seq can achieve sub-domain resolution in nucleolus

We first developed an analysis method to quantitatively characterize the layer structure of nucleolus, particularly the FC/DFC units. Using multi-color fluorescence immunostaining, we labeled each subdomain using corresponding marker proteins, i.e. the FC using RNA polymerase I subunit (RPA194), the DFC using fibrillarin (FBL) and the GC using nucleophosmin 1 (NPM1) in human foreskin fibroblast (HFF) cells (Fig. 1a). Unlike traditional ARTR-seq ^14^, all immunofluorescence experiments were performed on cells fixed with 4% PFA to preserve the tripartite structure of nucleoli. The GC layer of nucleoli can be easily distinguished from the FC/DFC unit using diffraction-limited fluorescence microscopy (Fig. 1b). However, the discrimination between the FC and DFC layers requires super-resolution microscopy. We applied single-molecule localization microscopy (SMLM) to two-color immunofluorescence labeled FC and DFC domains with CF568 dye and Alexa Fluor 647 (AF647) respectively (referred to as FC_AB_/DFC_AB_). The DFC signal formed a ring-like structure around the FC signal in the core (Fig. 1c and Supplementary Fig. 1a), as previously reported ^3^. As a comparison, in a sample where the DFC region was labeled by a mixture of two-color antibodies (referred to as DFC_AB_/DFC_AB_), both colors of the fluorescent signals exhibited the ring-like structure (Fig. 1c and Supplementary Fig. 1b).

We further applied correlation analysis to estimate the size of the FC/DFC unit (Extended Data Fig. 1, Supplementary Text). Importantly, the mean cross-correlation distance between the two colors (*D*_CF/AF_) in the FC_AB_/DFC_AB_ was 197 nm (median, 179 nm) (Extended Data Fig. 1c). This cross-correlation distance reflects the ring-like structure of DFC region surrounding the FC region and estimates the radius of the FC region, consistent with previously reported values ^3^. In contrast, the mean *D*_CF/AF_ in the DFC_AB_/DFC_AB_ was 11 nm (median, 0 nm), close to the resolution limit of SMLM imaging, suggesting that the two signals from the DFC region largely overlap (Extended Data Fig. 1c). In addition, autocorrelation analysis of each color allowed estimation of the ring thickness and the radius of the DFC region (Extended Data Fig. 1e-f). Collectively, these results from immunofluorescence samples demonstrate that SMLM imaging combined with correlation analysis is suitable for characterizing the spatial organization of nucleolar subdomains.

We next applied the above analytical framework to determine whether in situ RT provides sufficient spatial resolution for subdomain characterization. We analyzed the spatial relationship between the generated biotin-cDNA and the targeted protein in each nucleolar subdomain in HFF cells (Fig. 1a). Marker proteins of each layer were immunostained with CF568 dye, and the RT-generated biotinylated cDNA in each subdomain was stained with AF647-conjugated streptavidin. Using diffraction-limited microscopy, we observed that the biotin signal is much weaker when targeting the FC region compared to the DFC or GC (Fig. 1d). This observation is consistent with the enrichment of rDNA in the FC region and initiation of rDNA transcription at the FC-DFC interface ^3^. In addition, under diffraction-limited microscopy, FBL-targeted biotin signals were largely constrained within the FC/DFC units rather than diffusing into the GC region (Fig. 1d).

To better evaluate the spatial confinement of RT in the FC/DFC unit, we analyzed two samples in which biotin-cDNA generated from the DFC domain was imaged with either FC marker (FC_AB_/DFC_RT_) or DFC marker (DFC_AB_/DFC_RT_) using SMLM (Fig. 1e and Supplementary Fig. 1c-d) ^16,17^. In the FC_AB_/DFC_RT_ sample, AF647 signal from biotin-cDNA formed a ring-like structure surrounding the CF568 signal from the FC marker, suggesting that RT is spatially confined to the DFC region rather than penetrating into the FC domain. In contrast, in the DFC_AB_/DFC_RT_ sample, AF647 signal from biotin-cDNA formed the same ring-like structure as the CF568 signal from the DFC marker (Fig. 1e). We further employed correlation analysis to quantify the FC/DFC units (Extended Data Fig. 1g-h). In the FC_AB_/DFC_RT_ sample, the cross-correlation distance, *D*_CF/AF_, had a mean of 218 nm (median, 204 nm) (Fig. 1f), consistent with the radius of the FC region, as measured in the immunostaining control sample (Extended Data Fig. 1c). In contrast, *D*_CF/AF_ in DFC_AB_/DFC_RT_ had a mean value of 22 nm (median, 0 nm), again consistent with the immunostaining control sample (Figs. 1f and Extended Data Fig. 1c), supporting substantial overlap between biotin-cDNA and DFC marker signal. In addition, the thickness and radius of the biotin-cDNA from the DFC region, estimated by autocorrelation distance, were comparable to those of the DFC immunofluorescence signal (Figs. 1g-h and Extended Data Fig. 1e-f).

Collectively, these analyses confirm that the RT step in ARTR-seq is largely confined to the nucleolar subdomain, within the spatial resolution of SMLM.

### Differential enrichment of human transcriptome in nucleolar subdomains

We developed a pipeline to analyze nucleolar subdomain organization of human transcriptome (Fig. 2a). As the nucleolus is the primary site for ribosome biogenesis, reads were first mapped to annotated 47S rDNA reference sequence. Due to the repetitive nature of rDNA genes, we further depleted the residual 47S rDNA reads by mapping the reads to 18S, 28S and 5.8S rRNA sequences. The resulting 47S rDNA-depleted reads were then mapped to the entire human genome. Differential enrichment of non-47S rDNA genes among nucleolar subdomains was then calculated by DESeq2 analysis, relative to a control sample prepared identically except for the omission of primary antibodies (denoted as “–pri-AB control”).

**Fig. 2.**
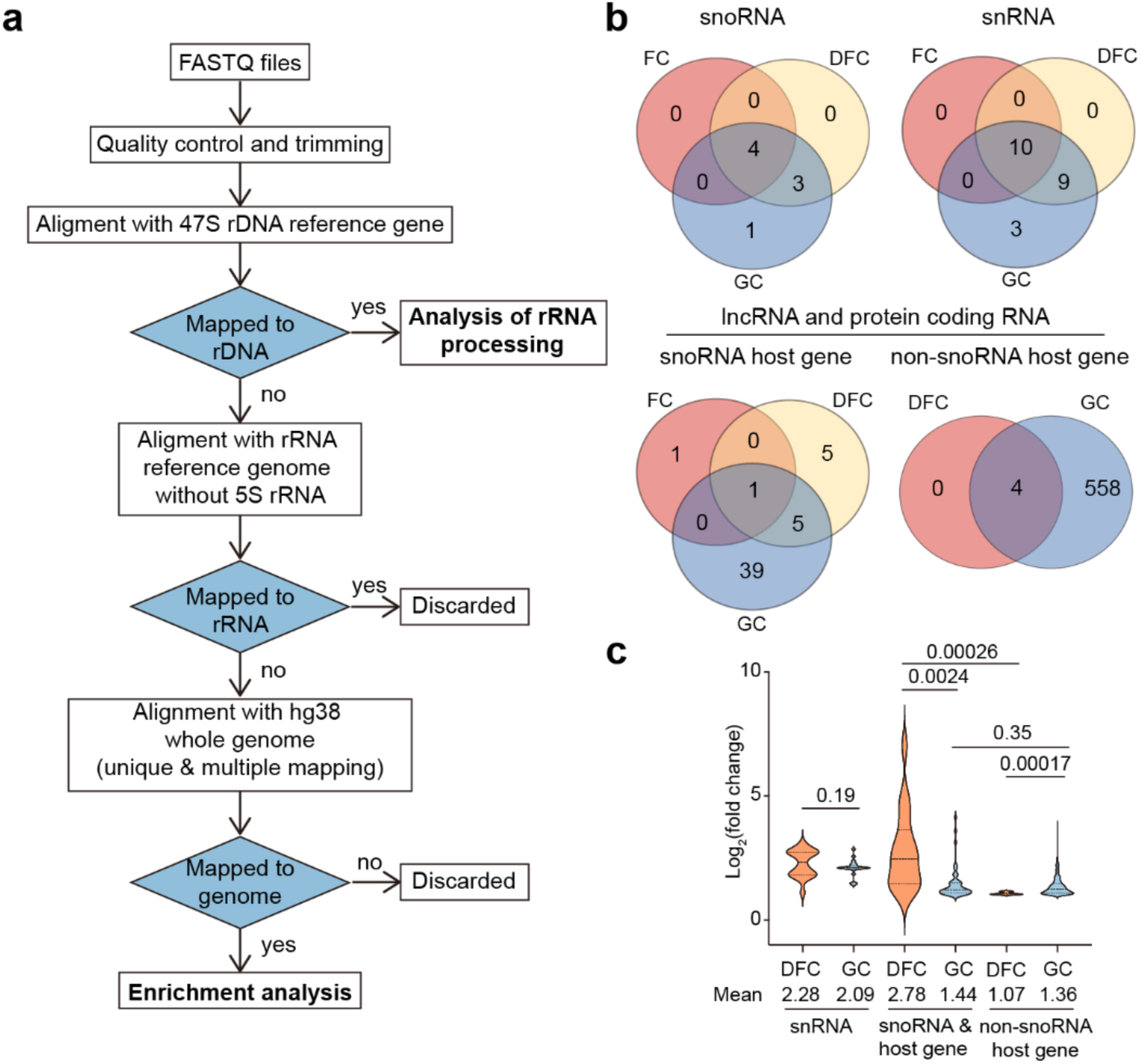
Transcriptome profiling of nucleolar subdomains. **a**, Workflow diagram of the sequencing data analysis for both rRNA and non-rRNA. **b**, Venn diagram of each transcript types enriched in three domains. Enriched genes were selected using criteria of log_2_(fold change) > 1, adjusted *p*-value < 0.05 and the standard error of log_2_(fold change) (lfcSE obtained from DESeq2) < 1. **c**, Violin plot shows the log_2_(fold change) values of (1) snRNA, (2) snoRNA and snoRNA host genes, and (3) non-snoRNA host genes in the DFC and GC. The protein-coding and lncRNA genes were divided into snoRNA host genes and non-snoRNA host genes according to snoDB database ^18^ (https://bioinfo-scottgroup.med.usherbrooke.ca/snoDB/). *P*-values were calculated by unpaired two-tailed t test.

Because many non-coding RNAs expected to localize to nucleoli, such as 5S rRNA and snoRNAs, are encoded by multiple gene copies, we considered reads that can only be mapped to a unique position (unique mapping) or mapped to multiple positions (multiple mapping). As expected, multiple mapping aligned significantly more reads to genes encoding 5S rRNA, snoRNAs, snRNAs and pseudogenes (Supplementary Fig. 2a) and identified more related genes accordingly (Supplementary Fig. 2b). The difference between using multiple and unique mapping was smaller for lncRNA and protein-coding genes (Supplementary Fig. 2a-b). The log_2_(fold change) calculated from unique mapping and multiple mapping showed a strong correlation across all subdomains (Supplementary Fig. 2c), suggesting that, for a subset of genes, the additional read counts assigned by multiple mapping are proportional between the two compared samples, resulting in consistent log_2_(fold change) values.

Applying criteria of log_2_(fold change) > 1, adjusted p-value < 0.05 and the standard error of log_2_(fold change) (lfcSE obtained from DESeq2) < 1, we compared the transcriptome, excluding 47S rDNA genes, within each nucleolar subdomain (Fig. 2b and Supplementary Table 1). Since most snoRNAs in human cells are encoded by snoRNA host genes, we further divided protein-coding and lncRNA genes into snoRNA host genes and non-snoRNA host genes. We chose to report snoRNA, snRNA, and snoRNA host genes using multiple mapping (excluding the corresponding pseudogenes), and genes encoding other transcript types, including non-snoRNA host protein-coding genes and lncRNAs, using unique mapping.

Transcripts from only a few non-rRNA genes were found to be enriched in the FC domain (Fig. 2b), including snoRNAs, snRNAs, and snoRNA host genes, which were mostly also enriched in the other two subdomains. The DFC domain was mostly enriched in snoRNA-related genes (Fig. 2b-c) ^18^, consistent with the canonical roles of snoRNAs in rRNA processing and modification ^19–21^. Several hundred genes were found enriched in the GC domain, including many non-snoRNA host protein-coding and lncRNA genes (Fig. 2b-c). Finally, 5S rRNA was found exclusively enriched in the GC domain and not enriched in the DFC, consistent with the spatial organization of 5S rRNA assembly to the ribosome ^22^. Collectively, our data reveal differential transcriptome organization across nucleolar subdomains. Below, we discuss the spatial organization of 47S rRNA, snoRNA, and non-snoRNA host protein coding and lncRNA transcripts individually.

### pre-rRNA processing demonstrates layered progression across nucleolar subdomains

We first asked whether spatial partitioning reflects functional progression of rRNA processing. In human cells, 47S pre-rRNA undergoes a series of cleavage and modification upon transcription to produce mature 18S, 28S and 5.8S rRNAs^4^, following a general temporal order: processing of 5’ ETS and 3’ ETS precedes ITS1 processing, which separates large and small subunit maturation, followed by ITS2 processing that separates 5.8S and 28S rRNA later in the pathway ^6,23^. Based on previous annotated endonuclease cleavage sites and pausing sites for exonuclease trimming ^23^, we divided the 47S rDNA sequence into various segments (Fig. 3a). We then normalized the reads mapped to each segment to the total reads mapped to entire 47S pre-rRNA sequence (Figs. 3b and Extended Data Fig. 2a). Finally, we calculated the enrichment index of each processing segment as the ratio of normalized reads in each subdomain to those of the –pri-AB control (Fig. 3c).

**Fig. 3.**
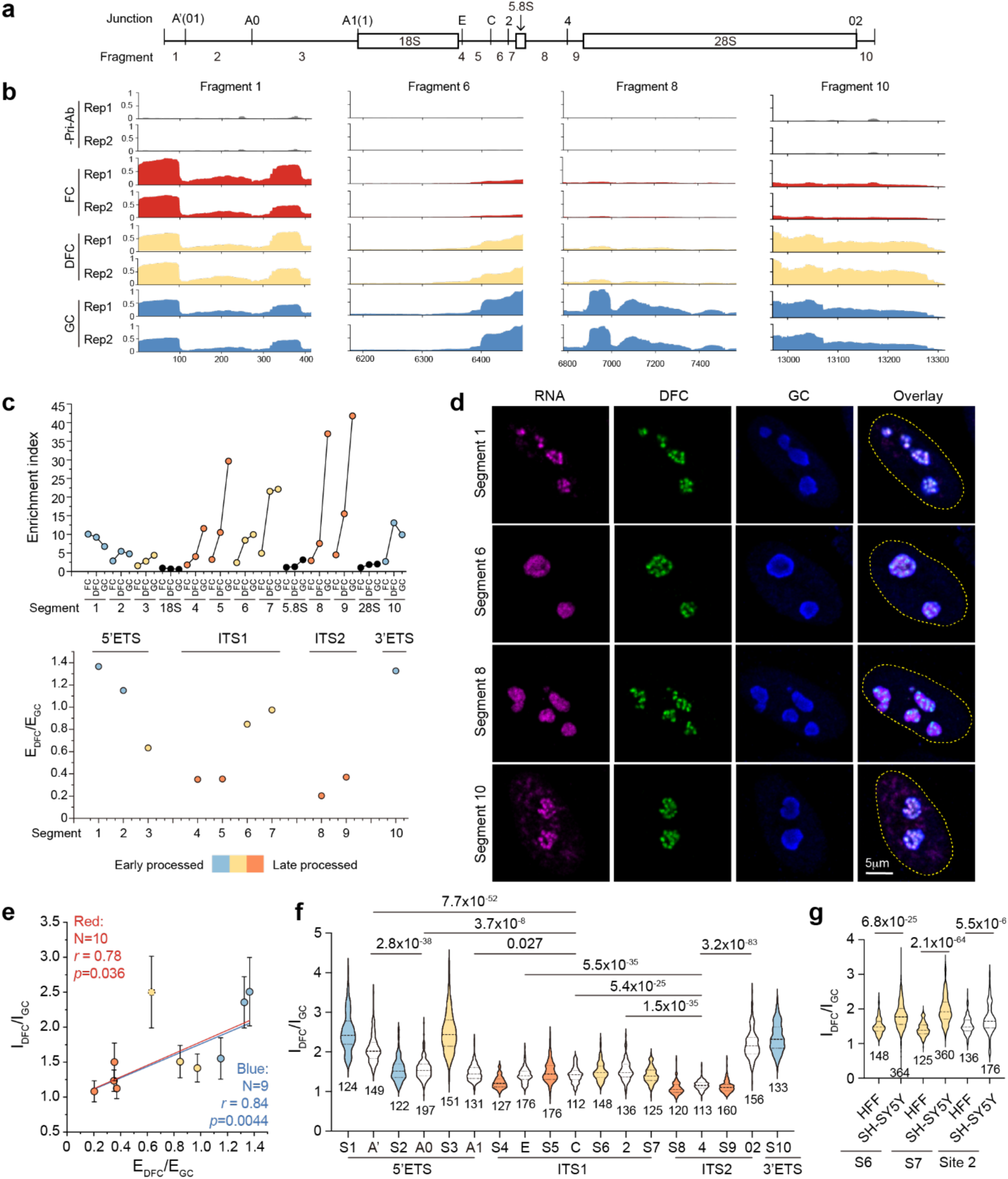
Distribution of rRNA processing intermediates in the nucleolar subdomains. **a**, Schematic representation of rRNA processing segments. **b**, Genome tracks of representative rRNA segments. For an easy visual comparison of the read distribution, the mapped reads were first normalized the total reads mapped to 47S the rDNA in each sample and then normalized to the maximum read across the four compared samples. **c**, Enrichment indices of different rRNA segments. The E_DFC_/E_GC_ in the lower panel presents ratio of enrichment index in the DFC to that in the GC. **d**, RNA FISH images of representative rRNA segments. Cell nuclei, stained with 4′,6-diamidino-2-phenylindole (DAPI), are outlined with dashed lines. **e**, Correlation between E_DFC_/E_GC_ from sequencing results and I_DFC_/I_GC_ representing the ratio of the average fluorescence intensities in the DFC to that in the GC from FISH images. Pearson’s correlation coefficient (*r*) and *p*-value for linear regression including all 10 segments (red line) or excluding segment 3 (shown in circle with dashed outline, blue line) are reported. **f**, **g**, I_DFC_/I_GC_ of rRNA fragments and junctions in FISH images. “S” denotes segment. “N” reports the number of segments in (e) and number of cells in (f), and (g), imaged from two biological replicates. *P*-values were calculated using two-sided t-test in (f) and (g).

Comparison of subdomain enrichment (E_FC_, E_DFC_ and E_GC_) of each segment revealed their spatial distributions (Fig. 3c). 5’ ETS and 3’ ETS segments were predominantly associated with high E_DFC_/E_GC_ ratios, indicating a high degree of DFC confinement. ITS1 contained segments with intermediate and low E_DFC_/E_GC_ values, indicating dispersion into the GC domain. Segments in ITS2 uniformly exhibited low E_DFC_/E_GC_ values, consistent with strong GC enrichment. The relative enrichment of each segment in the DFC compared to GC is consistent with the general temporal order during rRNA processing, i.e., 5’ and 3’ ETS are processed earliest, followed by ITS1 and then ITS2. This overall trend is consistent with two recent imaging-based studies ^24,25^. Within the ITS1 region, our data further revealed a higher E_DFC_/E_GC_ ratio for fragments 6 and 7, flanking the endonuclease cleavage site 2 (Fig. 3a), suggesting that this cleavage step, which separates the large and small subunit pre-rRNAs, and subsequent trimming of fragment 6 and 7 occur more in the DFC domain, whereases further processing towards the 3’ end of 18S rRNA occurs more in the GC domain.

To validate this sequencing result, we performed RNA FISH imaging on these segments (Figs. 3d and Extended Data Fig. 2b). Fluorescence intensity ratio between the DFC and GC domain (I_DFC_/I_GC_) showed a strong correlation with the E_DFC_/E_GC_ obtained from sequencing (Pearson’s *r* = 0.78, *p* = 0.036 for the red line, including segment 3, and Pearson’s *r* = 0.81, *p* = 0.0044 for blue line, excluding segment 3; Fig. 3e). However, the range of enrichment values differed between these two methods, as reflected by slopes of less than one in the linear fits. This difference likely reflects RT efficiency influenced by antibodies targeting different subdomains. Particularly, we noticed a discrepancy between sequencing and imaging results for segment 3. Segment 3 is confined within the DFC in imaging experiments but is slightly more enriched in GC in the sequencing results. This discrepancy may be related to its high GC content (up to 82%), which may affect the RT activity.

We further imaged the annotated junction regions between the segments (Extended Data Fig. 2c). The spatial distributions of these junction signals largely mirrored those of the adjacent segments (Fig. 3f). Specifically, endonuclease cleavage sites within 5’ and 3’ ETS were strongly enriched in the DFC, whereas junctions within ITS1 were intermediately DFC-enriched, and those within ITS2 are predominantly GC-localized. The overall consistency between the distributions of junctions and flanking segments indicates a strong spatial coupling between endonuclease cleavage events (leading to the loss of the junction signal) and exonuclease trimming events (leading to the loss of the segment signals).

Segment 3 again represents an exception, which showed a decoupled distribution from the flanking junctions (Fig. 3f). Within 5’ ETS, we observed that the A’ cleavage site was highly confined to the DFC domain, whereas the A0 and A1 sites were more dispersed to the GC domain, consistent with a recent imaging-based study ^25^, supporting that the endonuclease cleavage in the 5’ ETS follows a 5’-to-3’ direction. Interestingly, segment 3 between A0 and A1 remained highly DFC-confined despite the dispersion of the adjacent junctions ^24^. Given that segment 3 spans ∼2 kb, this observation indicates a possible finer spatial organization within this segment, where its internal region remains confined to the DFC, while the ends (close to A0 and A1 cleavage sites) extend into the GC domain (Extended Data Fig. 3a). Imaging with three sets of FISH probes targeting the 5’ end, middle and 3’ end region of segment 3 individually indeed supported such spatial organization (Extended Data Fig. 3a-c). This finer spatially organization of segment 3 may also contribute to the larger discrepancy between sequencing and imaging measurements (Fig. 3e).

Collectively, our data reveal a layered progression across nucleolar subdomains, which reflects the temporal order of rRNA processing. Particularly, we illustrate that the region between the A0 and A’ cleavage sites of the 5’ ETS exhibits internal organization across subdomains during processing.

### Differentiated cells exhibit more DFC-retained ITS1 processing

Ribosome biogenesis is tightly coupled to cell growth ^26^. Highly differentiated cells have lower ribosome synthesis output compared to highly proliferating cells, such as cancer cells ^27–29^. It was recently found that some differentiated cells exhibit 5′ ETS-centered small subunit processing impairments, which may contribute to a lower ribosome synthesis output ^24^. Specifically, in the neuroblastoma cell line SH-SY5Y, the position of segment 3 was observed to shift outward from the FC/DFC domain, with increased enrichment in the DFC and the peri-DFC domains ^24^.

We also applied our method to SH-SY5Y cells and found that, compared to HFF cells, segments 6 and 7 in SH-SY5Y cells showed markedly higher enrichment in the DFC, whereas the distributions of the other segments were largely consistent (Figs. 3c and Extended Data Fig. 4a). FISH imaging results of the corresponding segments confirmed this finding and in addition revealed a more DFC-retained junction signal corresponding to the endonuclease cleavage site 2 between segments 6 and 7 (Figs. 3a, 3g and Extended Data Fig. 4b). These results collectively suggest an increased DFC-retention of early steps during ITS1 processing in SH-SY5Y cells. This observation is consistent with an enlarged DFC domain compared to the GC domain in SH-SY5Y cells, despite an overall smaller nucleolar size (measured by the GC domain) compared to HFF cells (Extended Data Fig. 4c).

### snoRNAs are preferentially enriched in the DFC domain

We next examined the nucleolar enrichment of snoRNAs. Since snoRNA host genes typically encode multiple snoRNAs, we refined the snoRNA analysis using references for individually annotated snoRNA ^18^ (Supplementary Table 2). In HFF cells, the DFC domain contained the highest number of enriched snoRNAs (Fig. 4a), with the highest enrichment score (Fig. 4b). FC-enriched snoRNAs all overlapped with DFC- and GC-enriched snoRNAs, whereas GC-enriched ones all overlapped with DFC-enriched snoRNAs. These observations suggest that snoRNA localization peaks in the DFC domain, with minor fractions also distributed in FC and GC domains in HFF cells, consistent with their major function in rRNA modifications, which predominantly occurs in the DFC domain ^19,20^. We validated the observed trend by imaging snoRNAs generated from host genes GAS5 and SNHG29 using RNA FISH co-stained with markers for the DFC and GC domains (Figs. 4c and Extended Data Fig. 5-6). The RNA FISH signals confirmed enrichment in DFC over GC, as quantified by the intensity ratio (I_DFC_/I_GC_) (Fig. 4d).

**Fig. 4.**
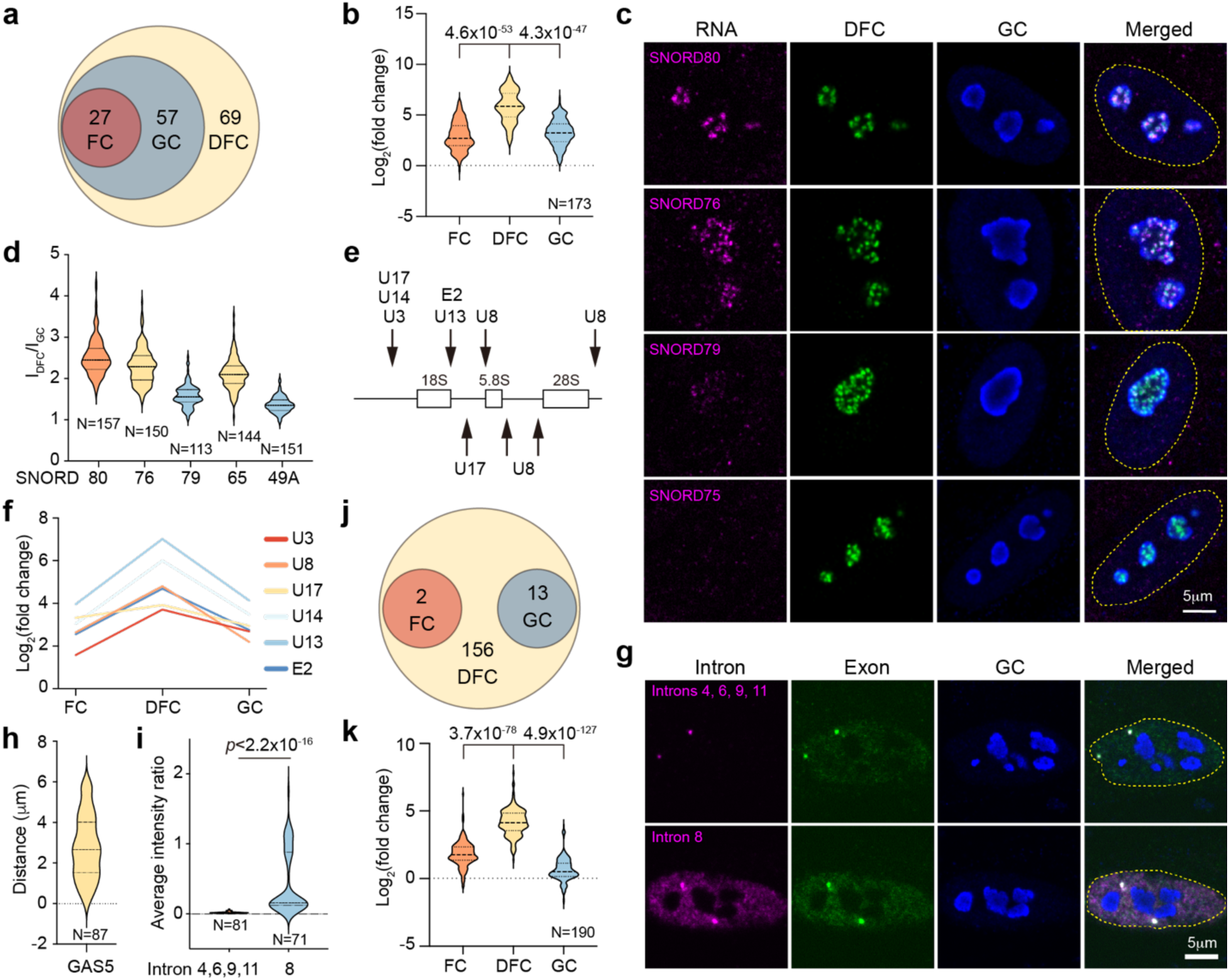
Identified snoRNAs show DFC-enriched distribution. **a**, Venn diagram of enriched snoRNAs in nucleolar subdomains of HFF cells. **b**, Violin plot showing the log_2_(fold change) values of snoRNA in the FC, DFC and GC of HFF cells. **c**, RNA FISH images of snoRNAs, co-stained with DFC and GC markers. Nuclei stained with DAPI are outlined by dashed lines. **d**, Ratios of fluorescence intensities within DFC over GC for imaged snoRNAs. **e**, Schematic diagram of rRNA processing steps regulated by snoRNAs. **f**, Violin plot showing the log_2_(fold change) values of snoRNAs marked in (e) across three subdomains. **g**, RNA FISH images of GAS5 introns 4, 6, 9, and 11 (top panel) and the SNORD80 surrounding regions of GAS5 intron 8 (bottom panel) together with exons. **h**, Violin plot of distance between GAS5 focus and the boundary of nucleolus. **i**, Ratio of average fluorescence intensity of RNA intron signal in nucleus to that at the transcription site (marked by bright foci using exon targeting probes). **j**, Venn diagram of enriched snoRNAs in nucleolar subdomains in SH-SY5Y cells. **k**, Violin plot showing the log_2_(fold change) values of snoRNAs in SH-SY5Y cells. Enriched snoRNAs were selected using criteria of total reads ≥ 10, log_2_(fold change) > 1 and adjusted *p*-value < 0.05. N reports number of genes in (b) and (k), and number of imaged cells in (d), (h), and (i), from two biological replicates. *P*-values were calculated using two-sided t-test.

SnoRNAs generally demonstrated robust enrichment in the DFC, reflected by the following observations. First, several snoRNAs reported to guide pre-rRNA processing, including U3 ^30^, U8 ^5,31,32^, U13 ^33^, U14 ^34^, U17 (E1) ^35,36^, and E2 ^37^ (Fig. 4e), also showed the highest enrichment in DFC (Fig. 4f), a subset of which have been consistently observed by imaging ^24^. As U13, U17 and E2 snoRNAs regulate ITS1 processing, and U8 is extensively involved in ITS2 processing, both of which occur in the GC domain, this observation suggests limited coupling between the spatial distributions of these snoRNAs and their guided rRNA processing segments (Fig. 3c). Second, snoRNAs containing different box types exhibited DFC enrichment except for AluACA RNAs. This class of snoRNAs, derived from primate-specific *Alu* repetitive elements, were not enriched in the nucleolus (Extended Data Fig. 7a), consistent with previous image analysis ^38^. Finally, snoRNAs with new annotated mRNA targets in addition to the rRNA targets ^39^ remained primarily enriched in nucleolus (Extended Data Fig. 7b).

To examine the spatial relationship between mature snoRNA and their precursor transcripts, we further analyzed the snoRNA host transcript GAS5 by RNA FISH using probes targeting mature snoRNA SNORD80, SNORD80-containing intron, additional introns reporting nascent transcripts, and exons of the host transcript (Extended Data Fig. 5). Signals from multiple introns appeared as discrete transcription foci distal from nucleoli (Fig. 4g-h), whereas exon signals showed colocalized foci with additional diffusive nucleoplasmic signal, indicating nuclear retention of the spliced GAS5 transcript (Fig. 4g). In contrast to the DFC-localized mature SNORD80 signal (Fig. 4d), the signal from the SNORD80-containing intron, in addition to the transcription sites, exhibited a uniform nucleoplasmic distribution excluding nucleoli (Fig. 4g, i). These observations suggest spatial segregation between snoRNA processing intermediates and mature snoRNAs.

Finally, we analyzed the snoRNA distributions in SH-SY5Y cells. Strikingly, although nucleolus-enriched snoRNAs largely overlapped between SH-SY5Y and HFF cells (Extended Data Fig. 7c), snoRNAs in SH-SY5Y cells were predominantly enriched in the DFC domain, with very few distributed into the FC or GC domains (Fig. 4j-k). This predominant DFC enrichment of snoRNAs is consistent with an increased DFC area (Extended Data Fig. 4c) and a higher degree of DFC retention in rRNA processing in SH-SY5Y cells (Fig. 3g).

### GC domain enriched non-snoRNA host transcripts are from nucleolar proximal genes

We examined the nucleolus-enriched non-snoRNA host transcripts, including protein-coding and lncRNA transcripts. These transcripts globally demonstrated low enrichment in GC domain (Figs. 2b and Supplementary Fig. 3). They contained high AT content, low GC content and long gene length primarily contributed long intron length (Fig. 5a-c), in contrast to nuclear speckle-enriched transcriptome ^13,15,40^. GC-enriched non-snoRNA host transcripts also contained reads mapped to both introns and exons (Fig. 5d) and exhibited a higher length-normalized intron fraction compared to a background consisting of all detected transcripts in this category (Fig. 5e), indicating that they represent unspliced pre-mRNA from nucleolar proximal genes.

**Fig. 5.**
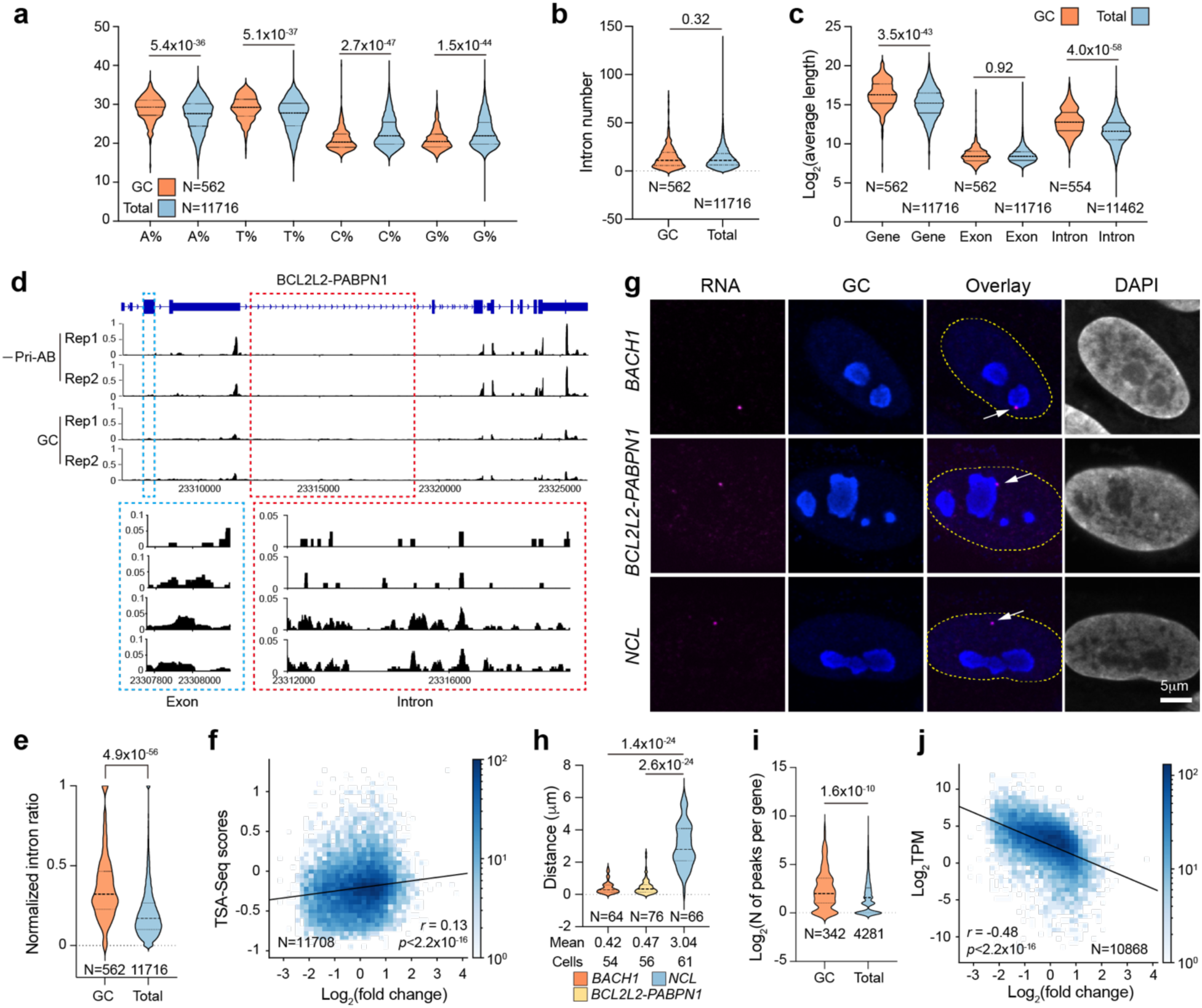
Features of GC-enriched transcripts. **a, b**, **c**, Violin plot of ATCG content (a), intron number (b), and gene length (c) of GC-enriched and total identified non-snoRNA host transcripts (lfcSE < 1). **d**, Genome track of a GC-enriched transcript *BCL2L2-PABPN1* with normalized reads. Dashed red and blue boxes show zoomed-in regions of an exon and intron respectively. For an easy visual comparison of the read distribution, the mapped reads were first normalized to the total rDNA-depleted reads against human genome in each sample and then normalized to the maximum read across the four compared samples. **e**, Violin plot of length-normalized intron fraction for GC-enriched and total detected non-snoRNA host transcripts. The reads mapped to the intron and exon of each gene were first normalized to the total intron and exon lengths of that gene respectively. Then the intron faction was calculated as the length-normalized intron reads divided by the sum of length-normalized intron and exon reads. **f**, Correlation between GC-domain enrichment index and TSA-Seq scores. **g**, Images of transcription sites of selected genes by RNA FISH (indicated with white arrows) and GC domain by immunofluorescence staining. Cell nuclei stained with DAPI are marked by the dashed lines. **h**, Violin plot of distance between each gene focus and the boundary of nucleolus. **i**, Violin plot of the number of H3K9me3 peaks per gene in GC-enriched and total detected non-snoRNA host transcripts. **j**, Correlation between enrichment index in the GC domain and poly(A) abundance of non-snoRNA host lncRNA and protein coding RNA. N reports number of genes in (a), (b), (c), (e), (f), (i) and (g), and number of cells in (h). Images were collected from two biological replicates in (h). *P*-values were all calculated using two-sided t-test.

To test this hypothesis, we compared the GC enrichment index of non-snoRNA host transcripts with their gene positions using published TSA-seq data for HFF cells ^41^. A weak positive correlation between GC enrichment index and TSA score suggests a modest association with gene position (Fig. 5f). To visualize the gene foci, we performed RNA FISH using probes against introns of two GC-enriched transcripts, *BACH1* and *BCL2L2-PABPN1*, along with a non-GC-enriched negative control, *NCL* (Fig. 5g). Compared to *NCL*, transcription sites corresponding to the gene foci of both *BACH1* and *BCL2L2-PABPN1* were located closer to the nucleoli (Fig. 5h), supporting the nucleolar proximity of their gene foci. Previous studies suggested that nucleolar-associated domains tend to be transcriptionally repressed and are enriched in repressive histone modifications, such as histone H3 lysine 9 trimethylation (H3K9me3) ^42–44^. Using published ChIP-seq data in HFF cells ^45^, we found that GC-enriched non-snoRNA host protein-coding and lncRNA genes contained significantly more H3K9me3 peaks per gene (Fig. 5i). Finally, higher GC enrichment correlated with lower poly(A) RNA abundance (Fig. 5j). Collectively, our data suggest that GC-enriched non-snoRNA host protein-coding and lncRNA transcripts are less spliced and tend to be generated from nucleolar-associated domains. These genes tend to contain more H3K9me3 markers and overall lower expression levels.

We noticed that *RPA194* transcript was specifically enriched in the RPA194-targeted dataset. While it is annotated as a putative snoRNA host gene, the reads were mapped exclusively to exon regions of the *RPA194* transcript rather than to the position of the annotated snoRNA (Extended Data Fig. 8a). We performed RNA FISH against exon regions of *RPA194* transcripts and found that they were uniformly distributed in the nucleus and cytoplasm (Extended Data Fig. 8b). Although the image result suggests that *RPA194* transcripts are not enriched in the FC domain, it raises the possibility that RPA194 protein may interact with its own mRNA.

## Discussion

In this study, we demonstrate that RNA species, like proteins, are spatially organized within layered MLOs at super-resolution. Using the nucleolus as a model system, ARTR-seq reveals distinct transcriptome partitioning across nucleolar subdomains and uncovers spatial organization of both rRNA processing intermediates and non-rRNA transcripts (Fig. 6).

**Fig. 6.**
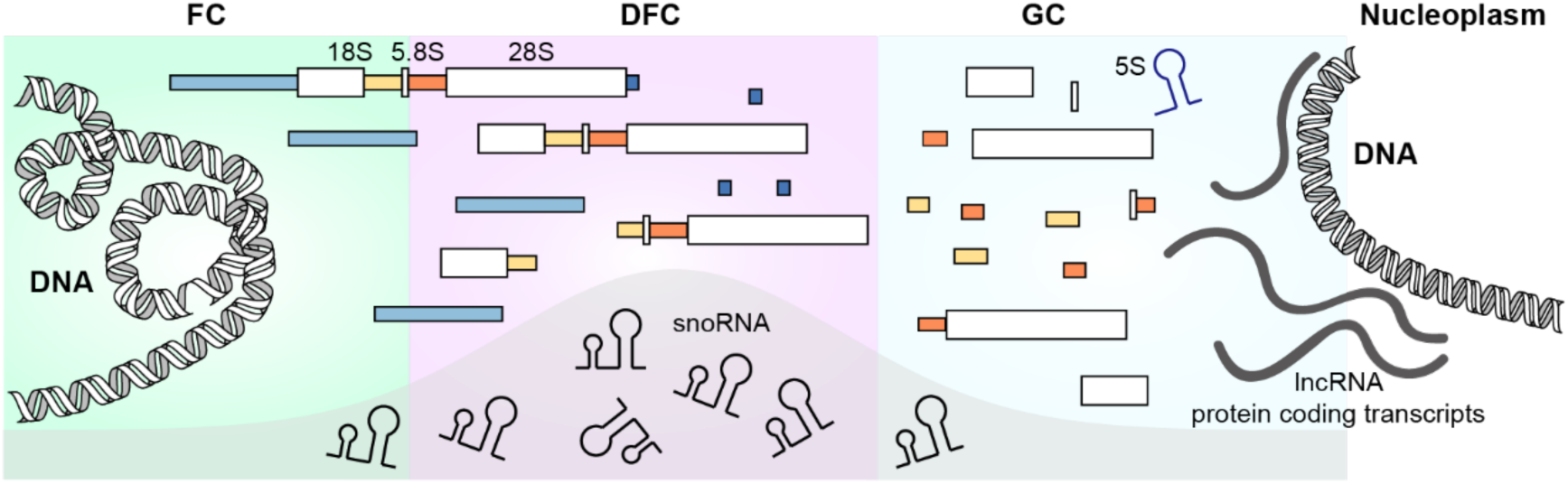
Organization of nucleolar transcriptome across different subdomains. Schematic illustration of the spatial organization of rRNA processing intermediates and non-rRNA transcripts, including DFC-enriched snoRNAs and GC-enriched transcripts from nucleolar-proximal genes.

Most prominently, this approach provides a spatially resolved view of rRNA maturation in a high-throughput way compared with imaging-based approaches and supports a layered progression of processing events across nucleolar subdomains ^24,25^. We find that processing of the 5′ ETS and 3′ ETS is largely confined to the DFC, early steps of ITS1 processing are more DFC-confined whereas later steps extend into the GC, and ITS2 processing occurs predominantly within the GC.

Interestingly, our data also indicate that a specific region of 5’ ETS exhibits internal organization across subdomain during processing. Compared to fast proliferating fibroblast cells, differentiated SH-SY5Y cells show an increased DFC-retention of ITS1 early processing steps that separates the small and large subunit maturation, consistent with a larger DFC size per nucleolus and reminiscent of a recent non-equilibrium model where rRNA processing impacts the FC patterning ^46^. Stalled ITS1processing has been observed as a mechanism to halt rRNA processing upon perturbations such as mTOR inhibition or nutrient deprivation ^47,48^. It is possible that the spatial retention of ITS1 processing in DFC might contribute to a slower outflow of rRNA processing and reduced ribosome synthesis output in differentiated cells.

We further find that the majority of C/D box and H/ACA box snoRNAs exhibit the highest enrichment in DFC, including those involved in both pre-rRNA modification and processing. Such spatial organization coincides with the primary pre-rRNA modification site ^21,49^ but has limited spatial correlation with their guided rRNA processing intermediates. This observation indicates that snoRNAs that guide pre-rRNA folding or cleavage may bind co-transcriptionally in the DFC domain as pre-rRNAs are transcribed from the FC/DFC interface. In contrast, non-snoRNA-related protein-coding and lncRNA transcripts display low to moderate enrichment in the GC domain. These transcripts are frequently incompletely spliced and tend to originate from nucleolus-proximal genes enriched in transcriptionally repressive histone modifications, consistent with their relatively low expression levels.

More broadly, this work establishes a framework for investigating the nanoscopic transcriptome organization within MLOs. The nucleolus provides a model system because its layered architecture is not only closely related to ribosome biogenesis, but also coordinates diverse nucleolar functions including genome organization, ribonucleoprotein assembly, and stress responses ^50–52^. Alterations in nucleolar morphology and architecture are associated with numerous human diseases ^53,54^. Furthermore, nucleolar localization of specific mRNAs has been observed during viral infection as part of immune regulatory strategies ^55^. Subdomain-resolved transcriptome profiling therefore provides a high-throughput approach to examine the organization of nucleolar activities across physiological or disease contexts. Higher-order structure is also a common feature of many MLOs. This strategy should therefore be broadly applicable to mapping transcriptomes within the subdomains of other biomolecular condensates.

Despite these advances, limitations should be considered. In situ RT depends on accessibility of RTase to target RNAs and may be affected by RNA secondary structure or by association with RNA-binding proteins. Consequently, detection efficiency for short and highly structured RNAs, such as certain snoRNAs, may be reduced. An unexpected observation was that ARTR-seq using RPA194 antibodies generated reads precisely mapped to all exons of *RPA194* mRNA, although imaging did not show clear enrichment of *RPA194* mRNA in the FC. One possible explanation is that RPA194 protein associates with its own mRNA, which may reflect previous observations that proteins tend to interact or condense with their cognate mRNAs ^56–60^. The potential functional significance of this interaction remains to be investigated.

## Methods

### Cell culture and treatment

HFF cells (ATCC, SCRC-1041) were cultured in Dulbecco’s modified Eagle’s medium (DMEM) supplemented with 15% (v/v) fetal bovine serum (FBS). SH-SY5Y cells (ATCC, CRL-2266) were cultured in a 1:1 mixture of DMEM and F12 medium supplemented with 10% (v/v) FBS. All mediums were supplemented with 100 U ml^−1^ penicillin and 100 µg ml^−1^ streptomycin, and cells were cultured at 37℃ in a 5% (v/v) CO_2_ atmosphere. Mycoplasma contamination was regularly tested in both cell lines. For fluorescence imaging and sequencing experiments, cells were seeded in an eight-well imaging chamber (no. 1.5 cover glass, Cellvis) at approximately 80% confluency.

### Reverse transcription in fixed cells

Unlike previously reported ARTR-seq ^14^, HFF or SH-SY5Y cells were fixed with 4% paraformaldehyde (PFA) for 10 min at room temperature to preserve tripartite structure of the nucleolus. The samples were then quenched with 125 mM glycine at room temperature and permeabilized with 0.5% Triton X-100 (Thermo Fisher Scientific) containing vanadyl ribonucleoside complexes (VRC, 2 mM, Sigma-Aldrich, #R3380) on ice. After that, samples were blocked with UltraPure bovine serum albumin (BSA) (1 mg ml^−1^, Thermo Fisher Scientific) for 0.5 h, stained with RPA194 (Santa Cruz, sc-48385, 1:100 dilution), fibrillarin (abcam, ab5821, 0.6 µg/mL), or NPM1 (Thermo Fisher Scientific, FC-61991, 1:50 dilution) antibodies at room temperature for 1 h, and then stained with secondary antibodies (mixture of donkey anti-rabbit and donkey anti-mouse IgG, Jackson ImmunoResearch, 1:400 dilution) at room temperature for 1 h. Next, samples were incubated with pAG-RTase for an additional 1 h to anchor reverse transcriptase to different subdomains. RNaseOUT (1 U μl^−1^, Thermo Fisher Scientific) was added at every step in the procedure above. A reverse transcription reaction mixture was prepared according to previous report with slight modification. 2 μM adapter-RT primer (5′-AGACGTGTGCTCTTCCGATCT-NNNNNNNNNN-3′), 0.05 mM biotin-16-dUTP (Jena Bioscience), 0.05 mM biotin-16-dCTP (Jena Bioscience), 0.05 mM dTTP (Thermo Fisher Scientific), 0.05 mM dCTP (Thermo Fisher Scientific), 0.1 mM dATP (Thermo Fisher Scientific), 0.1 mM dGTP (Thermo Fisher Scientific), and 1 U μl^−1^ RNaseOUT (Thermo Fisher Scientific) were mixed in 80 μl buffer of 50 mM Tris-HCl (pH 7.5) supplemented with 3 mM MgCl_2_ and 75 mM KCl. Cells were incubated in the reaction mixture at 37 °C for 1.5 h, followed by addition of 20 mM EDTA and 10 mM EGTA to terminate the reaction.

### Reverse transcription-based subdomain transcriptome sequencing

Following reverse transcription reaction, samples were digested with proteinase K (Thermo Fisher Scientific) at 37 °C overnight. The nucleic acids were recovered by phenol-chloroform extraction (pH 8.0), concentrated by ethanol precipitation, and digested by RNase H (1:80 dilution; NEB) and RNase A/T1 (1:20 dilution; Thermo Fisher Scientific) in RNase reaction buffer (40 μl, 50 mM Tris-HCl, pH 7.5, 75 mM KCl, 10 mM MgCl_2_, 10 mM DTT) at 37 °C for 1 h. The generated biotinylated cDNA was enriched by pre-blocked Dynabeads MyOne Streptavidin C1 (Thermo Fisher Scientific) at room temperature for 20 min. The 3′ cDNA adapter ligation was carried out by incubating the beads in the cDNA adapter ligation mixture (50 mM Tris-HCl, pH 7.5, 10 mM MgCl_2_, 25% PEG 8000, 1 mM ATP, 1 U μl^−1^ T4 RNA ligase 1 (NEB), and 5 μM of 3′ cDNA adapter (5′Phos-NNNNNNNNAGATCGGAAGAGCGTCGTGT-3′SpC3)) at 25 °C for 16 h. The biotinylated cDNA was eluted by elution buffer (95% (v/v) formamide in 10 mM EDTA (pH 8.0)) at 95 °C for 10 min), followed by ethanol precipitation. Libraries were generated using NEBNext Ultra II Q5 Master Mix (NEB) and purified on 6% Novex TBE gel (Thermo Fisher Scientific) with fragments size-selected between 180 and 400 bp. Next-generation sequencing was performed at the University of Chicago Genomics Facility on an Element Biosciences AVITI platform in single-end mode (150 bp).

### 47S rDNA mapping and rRNA processing analysis

FastQC was used to assess the quality of raw single-end FASTQ files. Adapter trimming was performed using Cutadapt (version 3.5). Reads were first aligned to the annotated 47S rDNA gene (GenBank: U13369.1) using STAR (v2.7.10b), allowing up to 30 multiple alignments per read (--outFilterMultimapNmax 30). Duplicate reads were removed using UMI-tools (v1.1.4) and quantified using the featureCounts ^61^ function in the Subread package (v3.0.5).

### Differential analysis of non-rRNA genes

Reads that were not mapped to rDNA gene above were further aligned to human rRNA sequences (28S, 18S and 5.8S) using STAR ^62^ (v2.7.10b) to further remove residual rRNA contamination except for 5S, which is not synthesized in nucleoli. Unmapped reads were then aligned to the human reference genome (GRCh38) annotated with GENCODE v39 using STAR (version 2.7.10b). Mapping was performed under two alignment stringency settings by varying the parameter --outFilterMultimapNmax (set to 20 or 1). Only reads with at least 24 matched bases were retained for subsequent analyses. Duplicates were removed with UMI-tools (v1.1.4) and read quantification was performed using the featureCounts function in the Subread package (version 3.0.5), generating read count values for each gene. Differential expression were carried out using DESeq2 ^63^ (v1.48.2) within the Bioconductor framework. For refined snoRNA analysis, 47S rDNA-depleted reads were aligned to annotated snoRNA genes ^18,22^ and quantified. The same differential analysis was then performed to calculate the enrichment of each snoRNA.

### Fluorescence labeling of FISH probes and secondary antibodies

For FISH probes labeling, oligonucleotides (Integrated DNA Technologies) were conjugated with amino–dideoxyuridine triphosphate (ddUTP) at the 3′ end by incubating 20 μl pooled oligonucleotides (100 μM), 6 μL Amino-11-ddUTP (1 mM; Lumiprobe, 15040), 1.2 μl terminal deoxynucleotidyl transferase (NEB, M0315L), and 3 μl 10x terminal transferase reaction buffer at 37 °C overnight. The products were purified by P-6 Micro Bio-Spin column (Bio-Rad) and subsequently reacted with 25 μg AF647 (Invitrogen) or CF568 (Sigma-Aldrich) NHS ester in 0.1M sodium bicarbonate (NaHCO_3_, pH 8.5) buffer at 37 °C overnight. Fluorescent probes were further purified by ethanol precipitation followed by P-6 Micro Bio-Spin column cleanup. The labeling efficiency was measured by UV-Vis spectroscopy. Probe sequences are listed in Table S3.

For secondary antibodies labeling, donkey anti-mouse IgG and donkey (715-005-150) anti-rabbit IgG (711-005-152) were purchased from Jackson ImmunoResearch. Mixture of antibodies (24 μl) were incubated with fluorophores (1 μg) in 1M NaHCO_3_ (pH8.5) buffer at room temperature for 1 h. Excess fluorophores were removed using P-6 Micro Bio-Spin column. The antibody-to-dye ratio was determined by UV-Vis spectroscopy.

### RNA FISH and immunostaining

Cells were fixed with 4% PFA (Electron Microscopy Sciences) at room temperature for 10 min, followed by permeabilization on ice in a solution containing 0.5% (v/v) Triton X-100 and VRC (2 mM) for 10 min. Samples were washed twice with 1× PBS, twice with 2× saline sodium citrate (SSC), and once with washing buffer (10% (v/v) formamide in 2× SSC). Hybridization buffer containing 10% (v/v) formamide, 10% (w/v) dextran sulfate, and 10 mM dithiothreitol (DTT) in 2× SSC was mixed with fluorescently labeled probe (5 nM each) and added to each well of the imaging chamber. Hybridization was carried out at 37 °C for 16 h in the dark. After that, cells were washed with washing buffer for 30 min at 37 °C to remove excess probes, followed by a second fixation with 4% PFA at room temperature for 10 min prior to immunostaining.

After three washes with PBS, the samples were blocked with blocking buffer (1 mg mL^−1^ BSA in PBS) at room temperature for 0.5 h. Primary and secondary antibody, diluted in blocking buffer at the concentrations described above, were sequentially incubated with the cells for 1 h each at room temperature. Cells were then washed three times with PBS, stained with 4′,6-diamidino-2-phenylindole (DAPI; Thermo Fisher Scientific, 62248), and stored in 4× SSC until imaging.

### Staining of biotin-cDNA in combination with immunostaining of marker proteins

For cases in which the biotin-cDNA signal and the antibody signal were from the same subdomain, such as DFC_AB_/DFC_RT_, the in situ reverse transcription was performed as described above, except that a fluorescently labeled secondary antibody was used during the immunostaining step prior to reverse transcription. After reverse transcription, the sample was blocked with BSA and incubated with AF647-labeled streptavidin (1 nM) at room temperature for 20 min. For cases in which the biotin-cDNA signal and the antibody signal were from different subdomains, such as FC_AB_/DFC_RT_, the reverse transcription was carried out as described above. After that, cells were washed twice with washing buffer (10% (v/v) formamide in 2×saline sodium citrate (SSC); 30 min followed by 15 min), post-fixed with 4% PFA, and blocked by UltraPure BSA (1 mg ml^−1^; Invitrogen, AM2618) for 30 min. Next, cells were immunostained with primary antibody (1 h), fluorescent secondary antibody (1 h; 1:200 dilution), and AF647-labeled streptavidin (1 nM; 20 min) at room temperature.

### Epi-fluorescence and confocal imaging

During imaging, cells were immersed in the imaging buffer containing Tris-HCl (50 mM, pH 8), 10% glucose, 2xSSC, glucose oxidase (0.5 mg ml^−1^, Sigma-Aldrich) and catalase (67 mg ml^−1^, Sigma-Aldrich). Diffraction limited epi-imaging was carried out on a Nikon TiE microscope equipped with a CFI HP TIRF objective (100×, NA 1.49; Nikon), an iXon Ultra 888 EMCCD camera (Andor), a 647 nm laser (Cobolt MLD), a 561 nm laser (Coherent Obis), and a 488 nm laser (Cobolt MLD). Confocal imaging was acquired on a Nikon Ti2-E inverted confocal microscope (Nikon AX-R) equipped with GaAsP PMT detectors (DUX-ST, Nikon) operating in resonant scanning mode and a CFI (Chromatic aberration-Free Infinity) Plan Apo objective (60× oil, numerical aperture (NA) 1.40, Nikon). Samples were excited using the AS405/488/561/640 laser unit (LUA-S4, Nikon) with appropriate laser and filter settings. Z-stacks (0.3 μm step size; five slices per stack) were taken for each channel with pinhole set to 1 Airy unit.

### Super-resolution imaging and image reconstruction

2D-SMLM was performed using the same microscope, objective, and EMCCD camera as in epi-fluorescence imaging, except total internal reflection fluorescence (TIRF) illumination was used. TetraSpeck microspheres (0.1 μm; Invitrogen) were diluted 1:500, added into each well, and incubated for 15 min at room temperature. Cells were washed with PBS and immersed in the imaging buffer containing 100 mM β-mercaptoethanol (BME; Sigma-Aldrich). For two-color SMLM, image sequences for the AF647 and CF568 channels were acquired sequentially using the JOBS module in NIS software. AF647 and CF568 were excited with 647-nm (40 mW) and 561-nm (85 mW) lasers, respectively. A 405-nm laser (CL2000; Crystal Laser) was used to stochastically activate fluorophores from the ‘off’ to the ‘on’ state. Image acquisition consisted of three consecutive frames of 647- or 561-nm excitation followed by one frame of 405-nm excitation, with an exposure time of 42 ms. The 405-nm laser power was gradually adjusted during imaging to maintain an appropriate density of molecules in the ‘on’ state; the maximum 405-nm power used in combination with the 647- and 561-nm lasers was 2.2 mW and 4 mW, respectively. A total of 9,000 frames were collected for each channel. Prior to SMLM imaging, an epi-fluorescence image containing at least one TetraSpeck bead within the region of interest was acquired for subsequent channel alignment.

ThunderSTORM ^64^ ImageJ plugin was used for SMLM image reconstruction. Molecules were first approximately localized using the ‘local maximum’ method with a peak intensity threshold set to 3 times the standard deviation of the residual background. Sub-pixel localization was then achieved using the integrated Gaussian PSF model with a fitting radius of 3 pixels (pixel size, 130 nm) and an initial sigma of 1.6 pixels. Connectivity was defined as an 8-neighborhood. Drift correction was performed using the cross-correlation method with a bin size of 20. Localizations with an xy-uncertainty greater than 40 nm were filtered out. Final images were rendered at 5× magnification.

### Correlation analysis

MATLAB was used for correlation analysis of FC/DFC units in SMLM images. For each two-channel region of interest (ROI) corresponding to each FC/DFC unit, 2D autocorrelation maps (CF568/CF568 channel and AF647/AF647 channel) and cross-correlation maps (CF568/AF647) were computed and normalized to correct for differences in the number of pixels involved in the correlation across displacements. The resulting 2D correlation maps were then radially averaged to obtain 1D radial correlation functions. A randomized control cross-correlation was generated by calculating the 1D radial cross-correlation between each channel and a pixel-shuffled version of the other channel; the average of these control correlations was subtracted from the measured radial cross-correlation to produce a background-subtracted radial correlation. Correlation length scales were quantified by fitting the 1D radial autocorrelation and background-subtracted cross-correlation curves to a single Gaussian or a double Gaussian function using nonlinear least-squares optimization. The resulting fit coefficients describing correlation amplitude, center, and width were extracted for each ROI for population average.

### Confocal image analysis

Nikon NIS-Elements software (AR 5.41.02), Fiji ImageJ2 and MATLAB was used for image analysis. Custom MATLAB codes were developed for image segmentation, enrichment analysis and distance calculation. Briefly, drift between channels was corrected and the most in-focus plane was subsequently selected via absolute Laplacian.

For GC and DFC enrichment analysis, fluorescence signals in DAPI, GC and DFC channels were first segmented using MATLAB’s binarize function individually. Incomplete cells at the image boundaries were removed from further analysis. Interactive user correction interface was implemented to allow manual correction in certain cases, such as splitting merged GC domains. Each segmented DFC domain was allocated to the corresponding GC domain, and each GC domain was allocated to corresponding nucleus by comparing overlapping pixels. Signals from RNA channel were then quantified in each segmented DFC, GC and nucleus domain. The relative RNA enrichment between DFC and GC domains for each cell was calculated by the ratio of mean RNA intensity in each domain.

For distance analysis between transcription sites and the GC domain, DAPI and GC channels were first segmented as described above. Transcription sites detected by RNA FISH were approximated using circular spot fitting with a radius of 5 pixels via manual annotation. Segmented GC domains and selected transcript sites were allocated into each corresponding cell (segmented DAPI region). The distance from each transcription site to the nearest GC domain was determined by the minimum distance between the center of the fitted spot and a pixel belonging to the boundary of a GC domain. A negative distance was assigned if the center of the transcription site fell within the GC domain.

## Supporting information

Table S2 snoRNA list

Table S1 gene list

Table S3 FISH probes

## Acknowledgements

We thank Dr. Xiaochang Zhang (The University of Chicago) for sharing the SH-SY5Y cells, Dr. Bei Lui (The University of Chicago) for useful discussion, and Dr. David Pincus (The University of Chicago) for feedback on the manuscript and Mr. Wei Liu for managing Fei Lab.

## Author contributions

Conceptualization: J.F., X.F.; Methodology and Investigation: J.F. (providing guidance for experimental approaches and analysis algorithms), X.F. (performing all experiments); Y.X. (providing pAG-RTase used for sequencing experiments). Funding acquisition: J.F., C.H; Project administration: J.F.; Supervision: J.F., W.-S.W.; Formal analysis: X.F., C.-C.T., J.H.M., S.-M.C., J.F.; Software: C.-C.T., J.H.M., I.T.L., J.F.; Writing – original draft: X.F., J.F.; Writing – review & editing: C.-C.T., J.H.M., I.T.L., Y.X., C.H.

## Data, code, and materials availability

Previously published H3K9me3 Chi-seq data are available under accession number GSE177895. All the sequencing data generated in this study and image analysis codes will be available upon acceptance of the manuscript.

---

**Extended Data Fig. 1.**
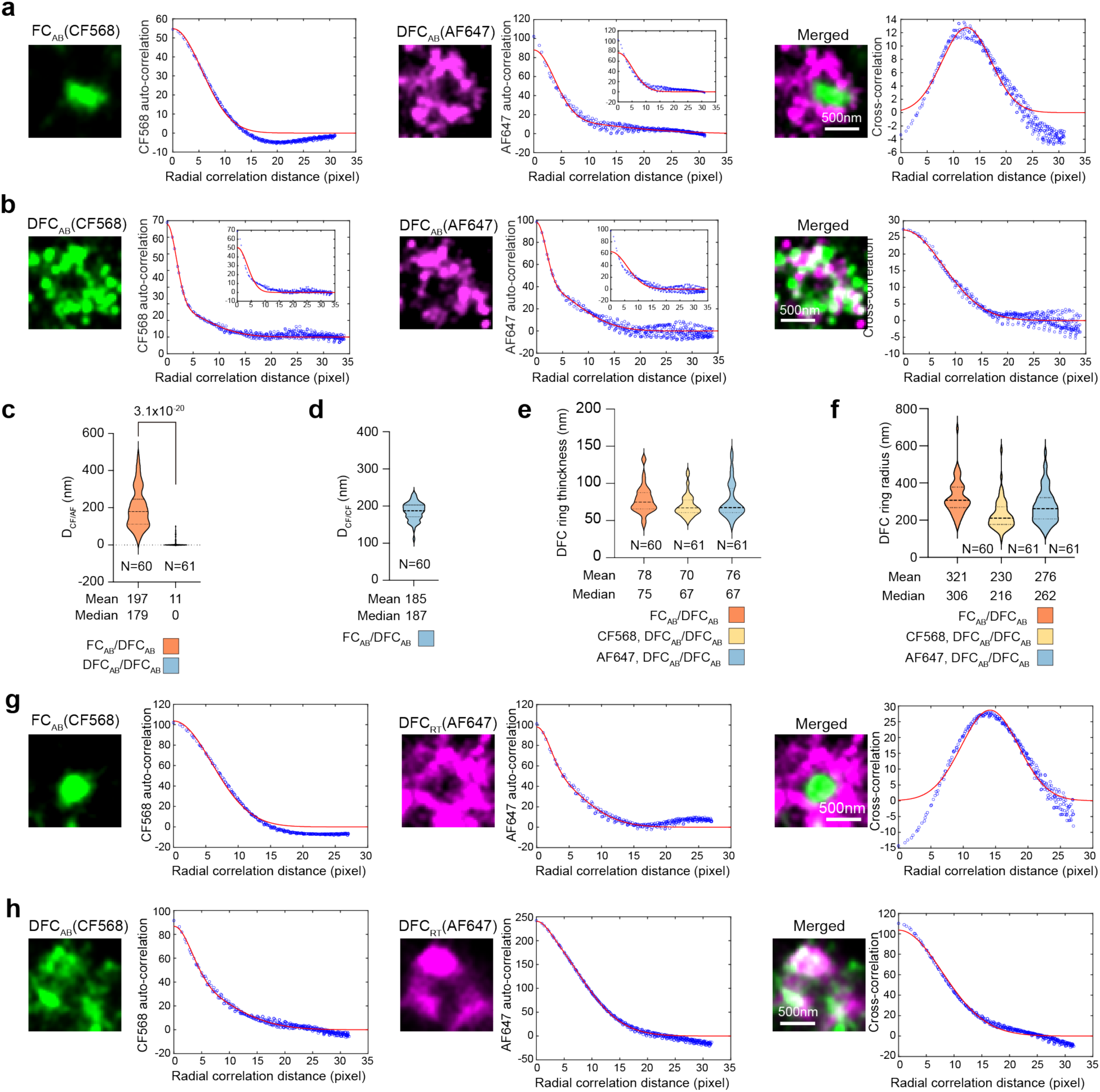
Correlation analysis of nucleolar structure. **a-b**, Autocorrelation analysis of individual color and cross-correlation analysis between two colors (Supplementary Text) for FC_AB_/DFC_AB_ (a) and DFC_AB_/DFC_AB_ (b) samples. The correlation distance for each FC/DFC unit was calculated by Gaussian fitting of the correlation curve. The autocorrelation curve of the DFC domain was better described using the two-peak Gaussian function. The one-peak Gaussian function is reported as an inset plot for comparison. **c-f**, Violin plot for (c) cross-correlation distances between AF647 and CF568 channels of the two samples, *D*_CF/AF_; (d) The radius of the FC domain, estimated by autocorrelation distance, *D*_CF/CF, 1_; (e) The ring thickness of the DFC domain, estimated by autocorrelation distance, *D*_AF/AF, 1_ or *D*_CF/CF, 1_; (f) The radius of the DFC domain, estimated by autocorrelation distance, *D*_AF/AF, 2_ or *D*_CF/CF, 2_. The distance was calculated using the fitted correlation distance times the effective pixel size (26 nm) in our SMLM imaging. **g-h**, Autocorrelation analysis of individual color and cross-correlation analysis between two colors for FC_AB_/DFC_RT_ (g) and DFC_AB_/DFC_RT_ (h) samples. The correlation distance for each FC/DFC unit was calculated by Gaussian fitting of the correlation curve. The autocorrelation curve of the DFC domain was also better described using the two-peak Gaussian function. N in (c)-(f) reports the number of FC/DFC units in each plot, collected from 5-6 cells from two biological replicates. *P*-values were calculated using two-sided t-test.

**Extended Data Fig. 2.**
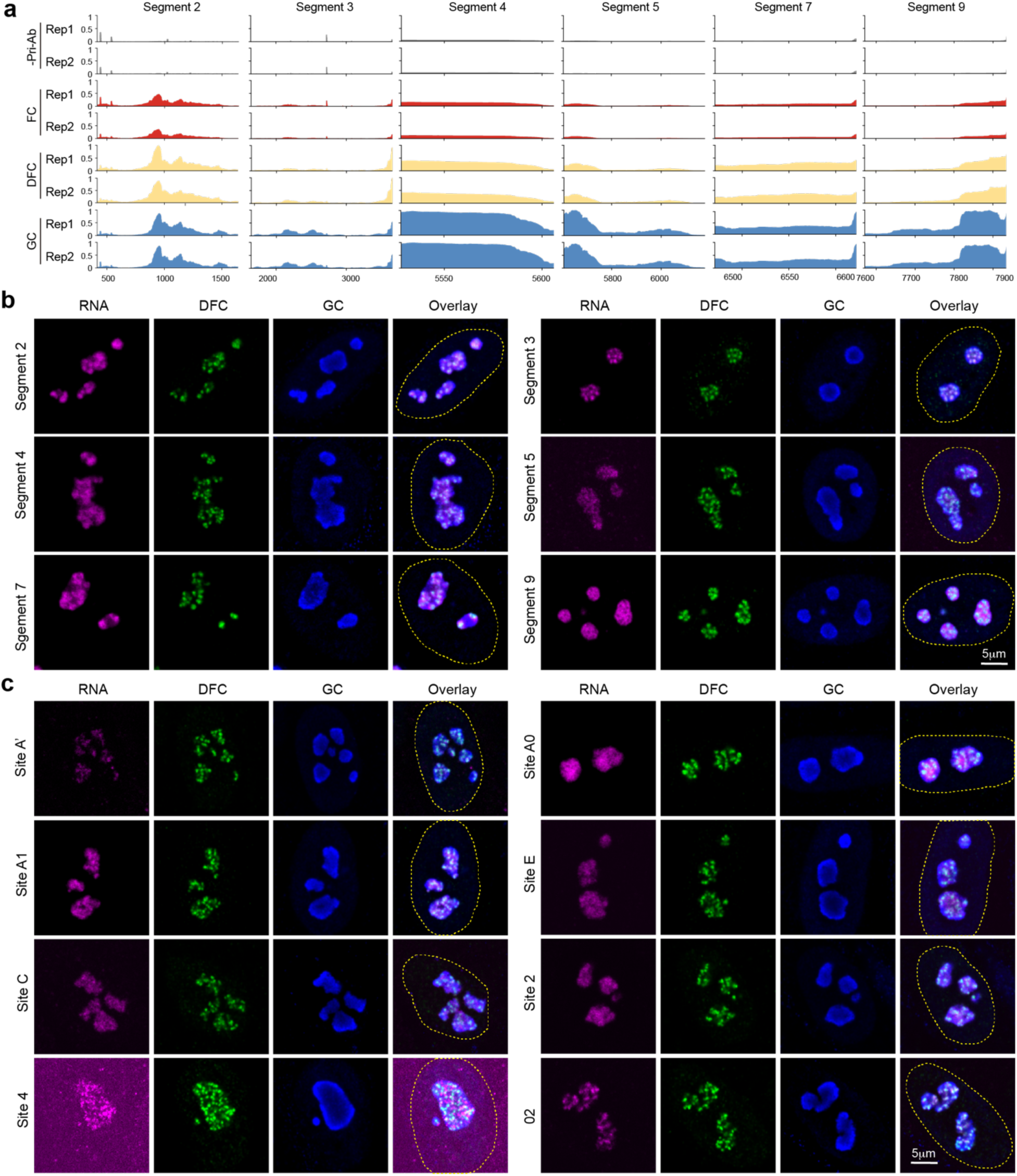
Distribution of additional rRNA processing intermediates in the nucleolar subdomains. **a**, Genome tracks of rRNA segments other than those shown in Fig. 3b. **b**. RNA FISH images of rRNA fragments and junctions in HFF cells. Cell nuclei were outlined by yellow dashed lines.

**Extended Data Fig. 3.**
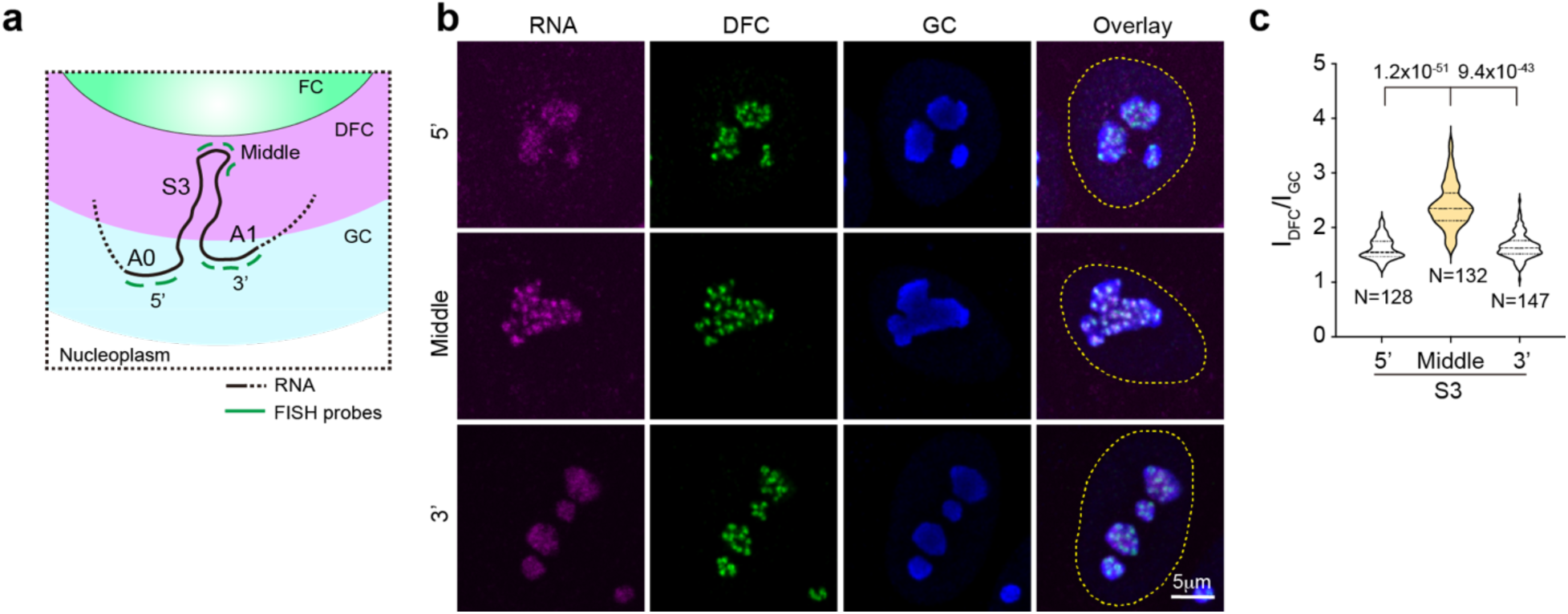
Organization of rRNA segment 3 in nucleolus. **a**, Schematic model of segment 3 organization in nucleolus. Green bars marked the FISH probe targeting regions in the 5’ end, middle, and 3’ end of segment 3. **b**, RNA FISH images using the three sets of probes illustrated in (a). **c**, I_GC_/I_DFC_ of different regions of rRNA segment 3 in FISH images. N denotes the number of cells imaged in two biological replicates. *P*-values were calculated by two-sided t-test.

**Extended Data Fig. 4.**
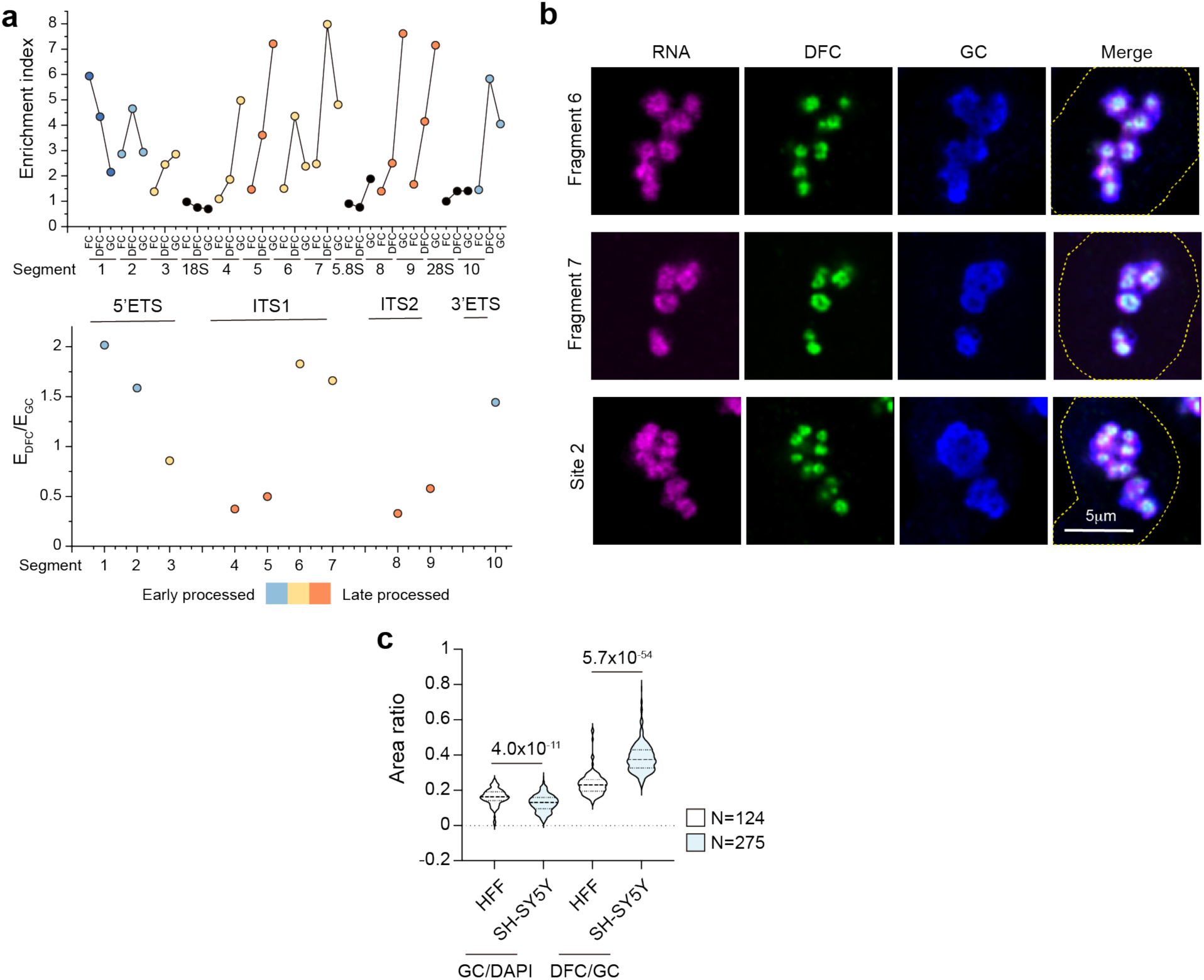
Nucleolar distribution of rRNA processing intermediates in SH-SY5Y cells. **a**, Enrichment indices and E_DFC_/E_GC_ of each fragment in the nucleolar subdomain. **b**, RNA FISH images of rRNA segments 6 and 7 and junction corresponding to cleavage site 2, co-stained with the DFC and GC marker proteins in SH-SY5Y cells. Cell nuclei were outlined by dashed lines. **c**, Area ratios of GC to nucleus (quantified by DAPI signal) and of DFC to GC in HFF and SH-SY5Y cells. N reports the number of cells imaged in two biological replicates. *P*-values were calculated by two-sided t-test.

**Extended Data Fig. 5.**
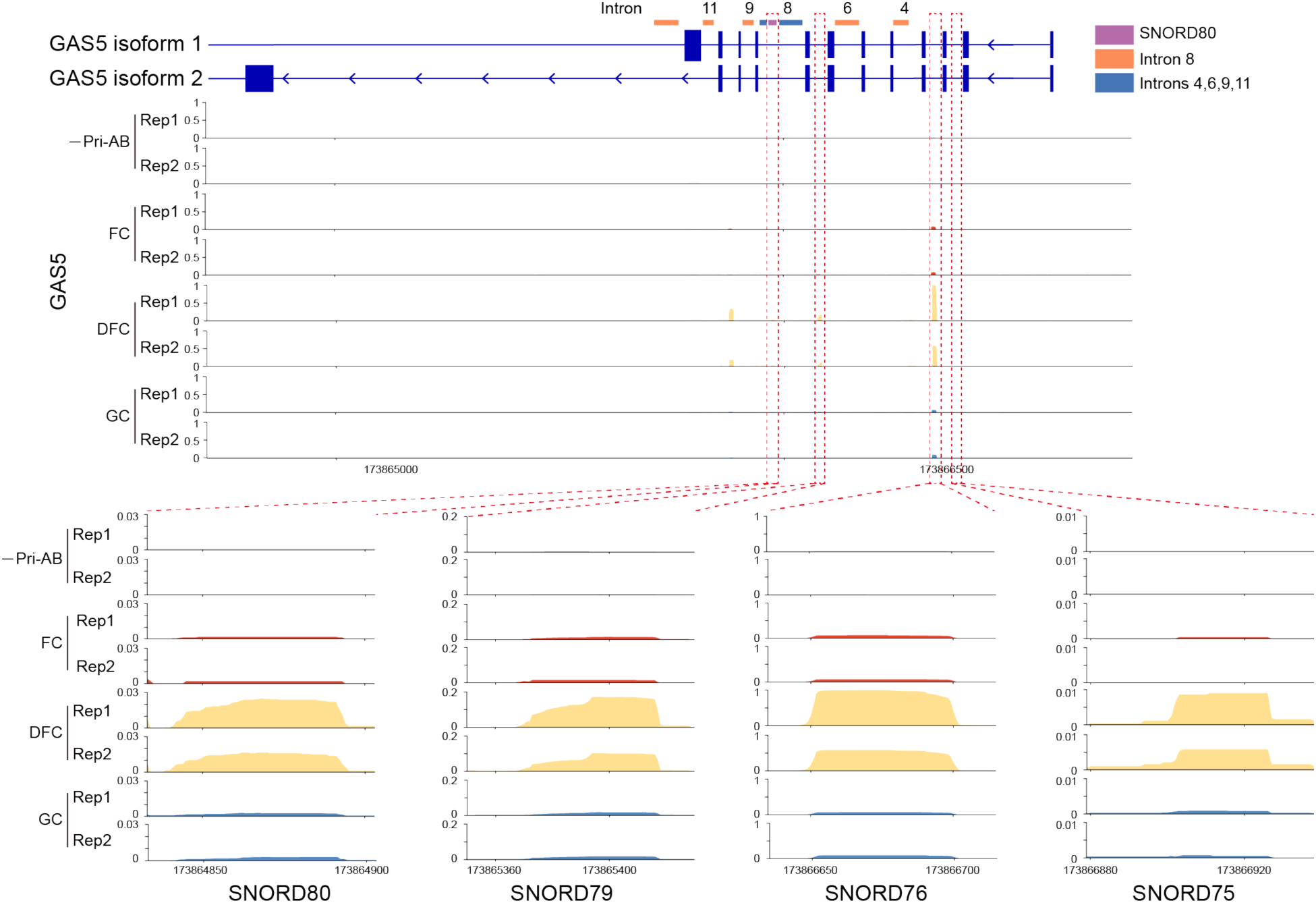
Genome track plots of GAS5 with zoomed-in regions of selected encoded snoRNAs. Colored lines depict FISH probes targeting regions.

**Extended Data Fig. 6.**
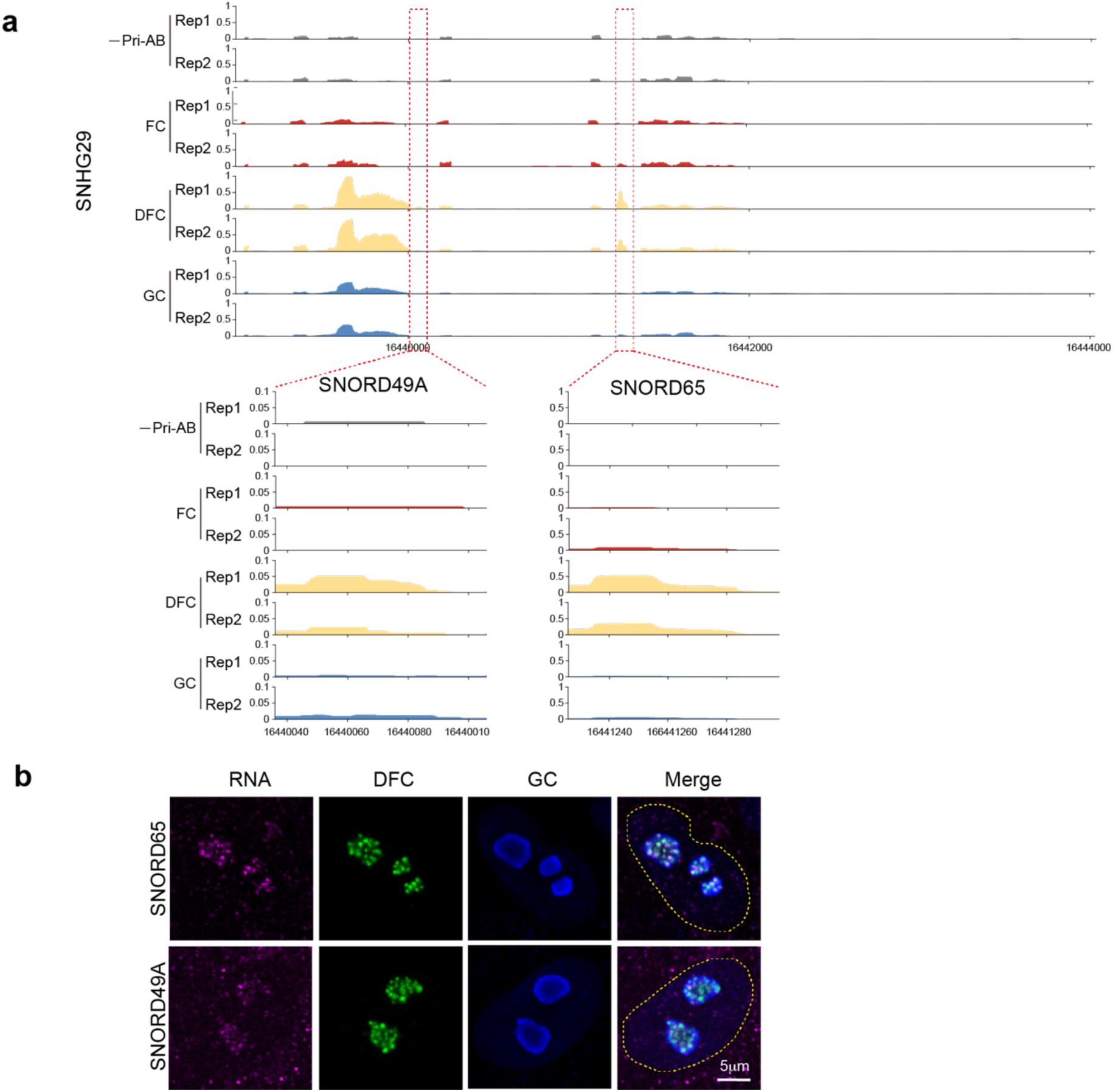
Localization of snoRNAs from host gene SNHG29. **a**, Genome tracks showing the enrichment of selected snoRNAs from SNHG29. **b**, RNA FISH imaging for snoRNAs from SNHG29, co-stained with the DFC and GC domain marker proteins.

**Extended Data Fig. 7.**
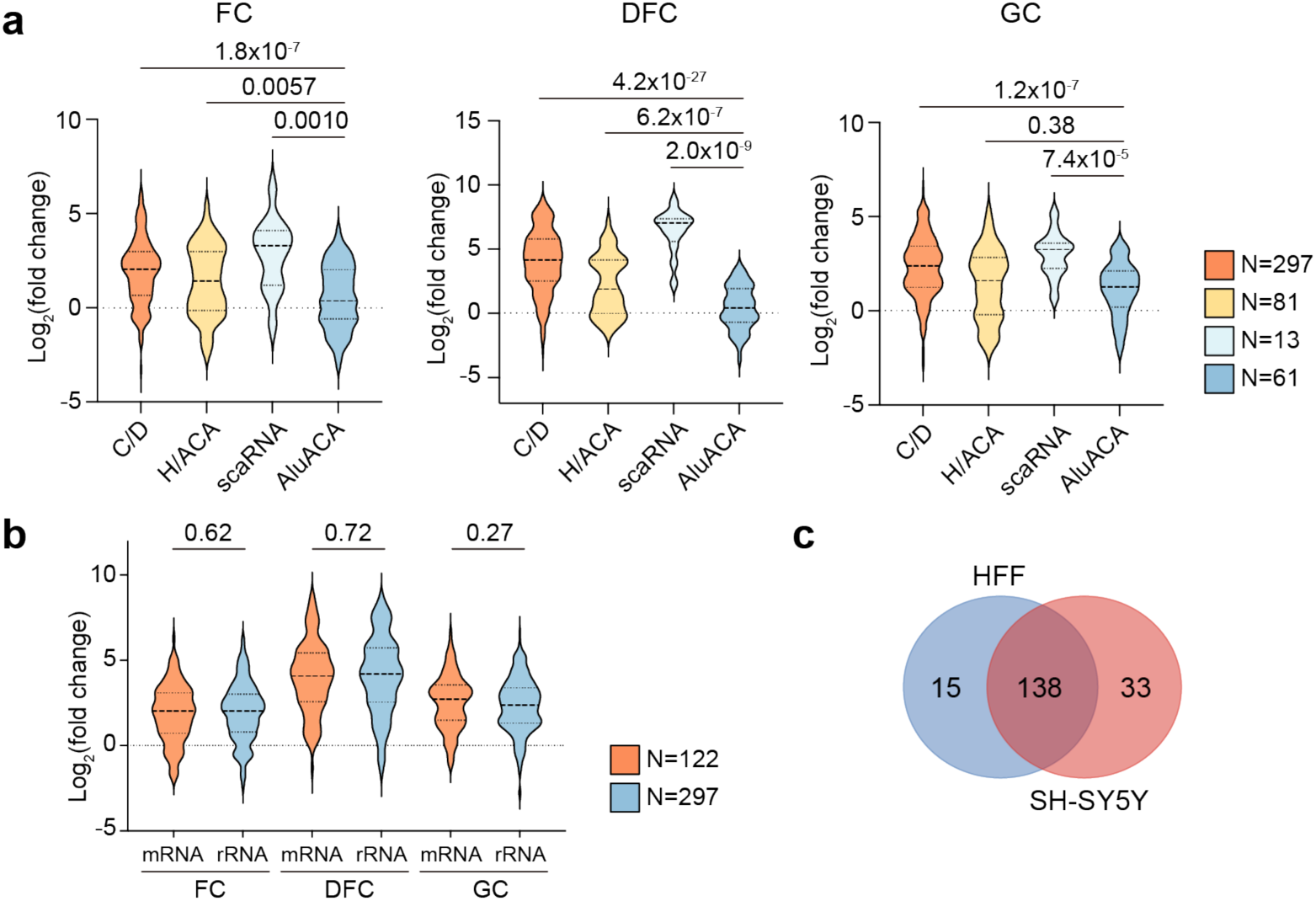
Enrichment of snoRNAs in HFF and SH-SY5Y cells. **a**, Violin plots of log_2_(fold change) values in each subdomain for snoRNAs grouped by box type. **b**, Violin plots of log_2_(fold change) values in each subdomain for snoRNAs categorized by their target RNAs. Total mRNA-targeting snoRNAs from all five tested human cell lines were included from reference ^39^. SnoRNAs with total reads ≥ 10 were included for plotting, Gene numbers are reported by N. *P*-values were calculated using two-sided t-test. **c**, Venn diagram of nucleolus-enriched snoRNAs in HFF and SH-SY5Y cells. Enriched snoRNAs are selected based on criteria: total reads ≥ 10, log_2_(fold change) > 1 and adjusted *p*-value < 0.05.

**Extended Data Fig. 8.**
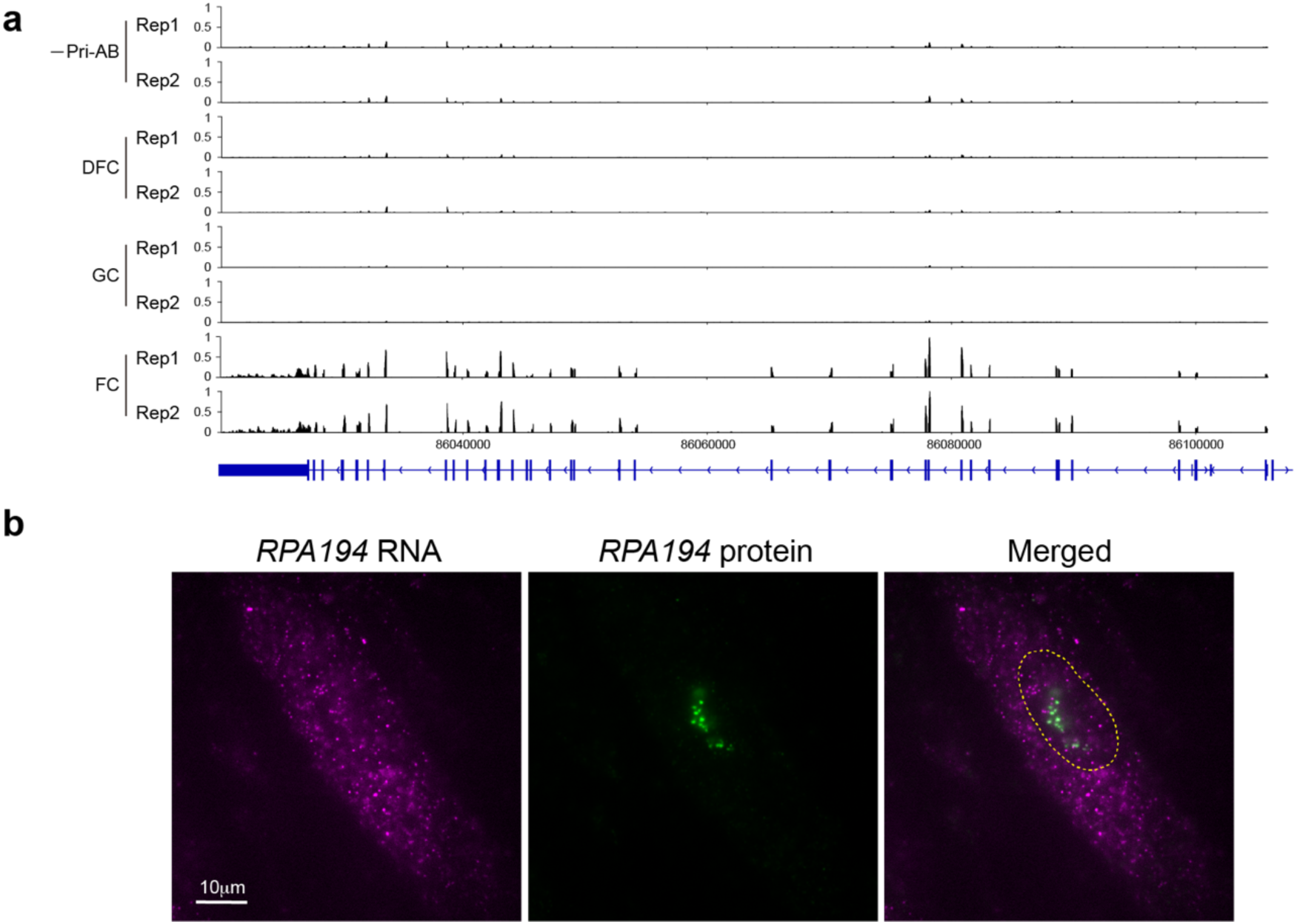
*RPA194* are not enriched in the FC domain. **a**, Genome tracks of reads mapped to PRA194 gene via targeting PRA194 protein. Reads were exclusively aligned to the exons. **b**, RNA FISH images of *RPA194* mRNA co-stained with antibodies against PRA194 protein. Nuclei are outlined by dashed lines.

