## Supplementary information for "Super-resolved spatial organization of the nucleolar transcriptome"

Xinqi Fan et al

**The PDF file includes:**

Supplementary Text

Supplementary Fig. 1-3

References

**Other Supplementary Material for this manuscript includes the following:**

Supplementary Tables 1-3

### Supplementary Text

#### *Correlation analysis of SMLM images of FC/DFC unit*

To quantify the dimensions of the FC and DFC layer from the $FC_{AB}/DFC_{AB}$ image, we employed cross-correlation and autocorrelation analysis. Cross-correlation versus radial distance profile of CF568 and AF647 signals from individual FC/DFC unit exhibited a peak centered at a non-zero position (Extended Data Fig. 1a). The cross-correlation versus radial distance profile was fit with a Gaussian function, and the center of the Gaussian reports the cross-correlation distance between CF568 and AF647 ($D_{CF/AF}$) for each FC/DFC unit. Analysis of tens of FC/DFC units yielded a mean $D_{CF/AF}$ of 197 nm, with a median of 179 nm (Extended Data Fig. 1c). This cross-correlation distance is consistent with the ring-like structure of the DFC region surrounding the FC region and estimates the average radius of the FC region (Extended Data Fig. 1c) [1].

The autocorrelation of CF568 signal from the FC region and the AF647 signal from the DFC region of the $FC_{AB}/DFC_{AB}$ sample both showed an autocorrelation versus radial distance profile peaked at zero, as expected. A single Gaussian function adequately described the CF568 autocorrelation versus radial distance profile (Extended Data Fig. 1a). The width of the Gaussian function reports the autocorrelation distance ($D_{CF/CF}$), yielding a mean value of 185 nm and median of 187 nm (Extended Data Fig. 1d). $D_{CF/CF}$ should also reflect the radius of the FC region and is indeed in consistency with the above estimated $D_{CF/AF}$ value. The AF647 autocorrelation versus radial distance profile, however, cannot be described well by a single Gaussian function, but was well described by a double Gaussian function with different widths (Extended Data Fig. 1a). We reasoned that the shorter autocorrelation distance ($D_{AF/AF,\,1}$), defined by the width of the narrower, higher Gaussian peak, reflects the thickness of the DFC ring, and the longer autocorrelation distance ($D_{AF/AF,\,2}$), defined by the broader, lower Gaussian peak, reflects the overall size of the DFC ring (Extended Data Fig. 1a). $D_{AF/AF,\,1}$ had a mean value of 78 nm with a median value of 75 nm, consistent with the thickness of DFC ring reported in previous super-resolution imaging [1] (Extended Data Fig. 1e). $D_{AF/AF,\,2}$ had a mean value of 321 nm with a median value of 306 nm, consistent with the overall dimension of the FC/DFC unit (Extended Data Fig. 1f).

As a control, in the $DFC_{AB}/DFC_{AB}$ image (Extended Data Fig. 1b), $D_{CF/AF}$ had a mean value of 11 nm and a median value of zero, suggesting that CF568 and AF647 signals both from the DFC region largely overlapped (Extended Data Fig. 1b, c). Consistent with the AF647 signal in the $FC_{AB}/DFC_{AB}$ image, both CF568 and AF647 signals in the $DFC_{AB}/DFC_{AB}$ image required double Gaussian fitting of the autocorrelation versus radial distance profile, confirming that the presence of the second longer correlation distance is not due to the use of different fluorophores, but reflects the geometry of the DFC ring (Extended Data Fig. 1b). The thickness of the DFC ring and the overall size of the FC/DFC unit estimated from both CF568 and AF647 in the $DFC_{AB}/DFC_{AB}$ image were consistent with that estimated from the AF647 signal in the $FC_{AB}/DFC_{AB}$ image above (Extended Data Fig. 1e-f).

It should be mentioned that the CF568 and AF647 signals from the DFC region are not expected to completely overlap even though they label the same DFC layer. Several factors could contribute to this observation, including (1) incomplete immunostaining efficiency and mutual exclusion of differently labeled antibodies from binding to the same molecule, (2) incomplete sampling of fluorophores inherent to SMLM imaging, where sparse activation is required for single-molecule detection, and (3) the possible uneven distribution of FBL proteins in the DFC layer.

The same correlation analysis was applied to $FC_{AB}/DFC_{RT}$ and $DFC_{AB}/DFC_{RT}$ samples.

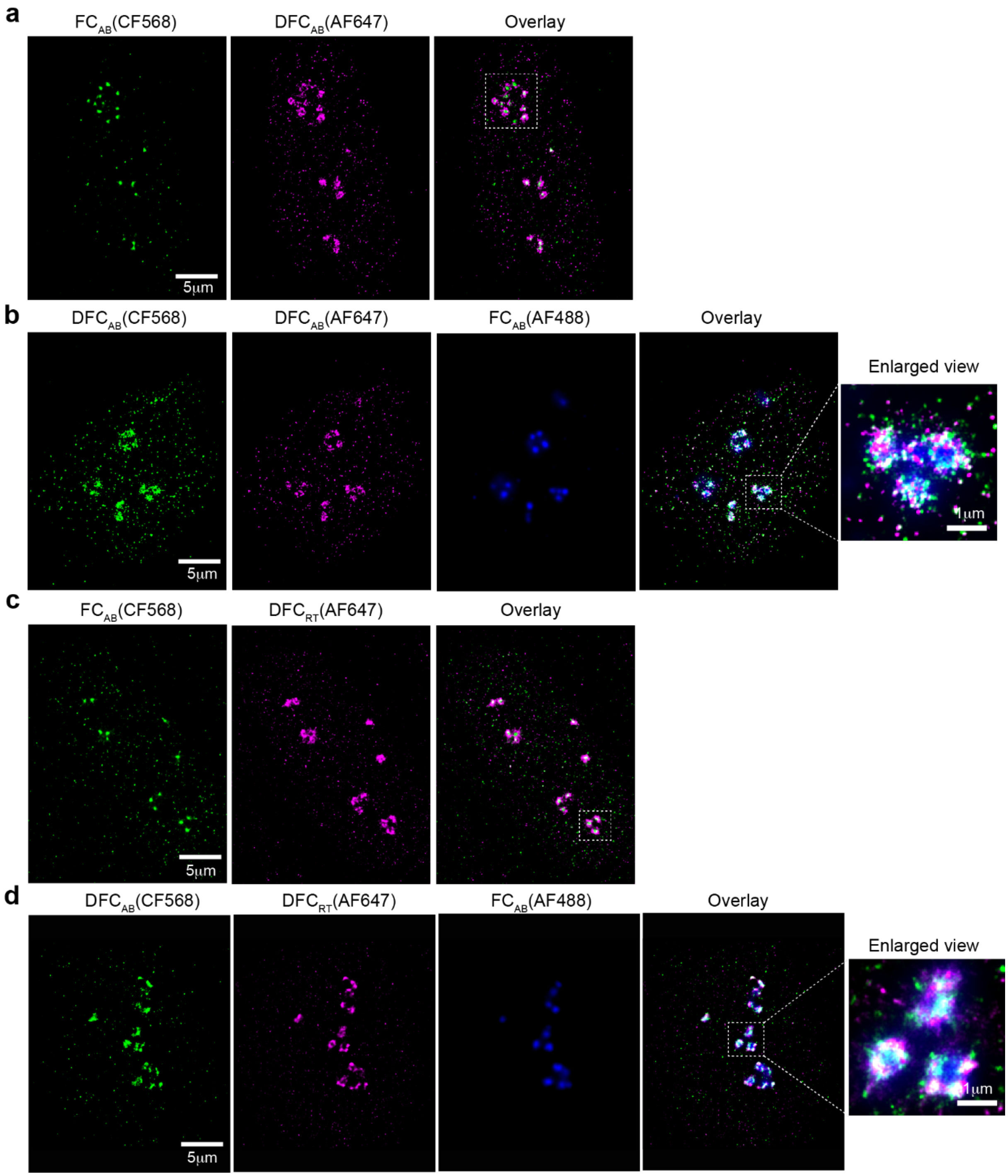


**Supplementary Fig. 1 | Uncropped images of FC/DFC units of nucleoli within a single cell.** **a**, **b**, Immunofluorescence images of FC/DFC units in a single HFF cell recorded by SMLM. Boxed regions are enlarged in Fig 1c. **c, d**, SMLM images of biotin-cDNA generated in DFC domain together with immunofluorescence staining of FC or DFC domain a single cell. Boxed regions are presented in Fig 1e. FC channels in (b) and (d) were recorded by epi-fluorescence microscopy.

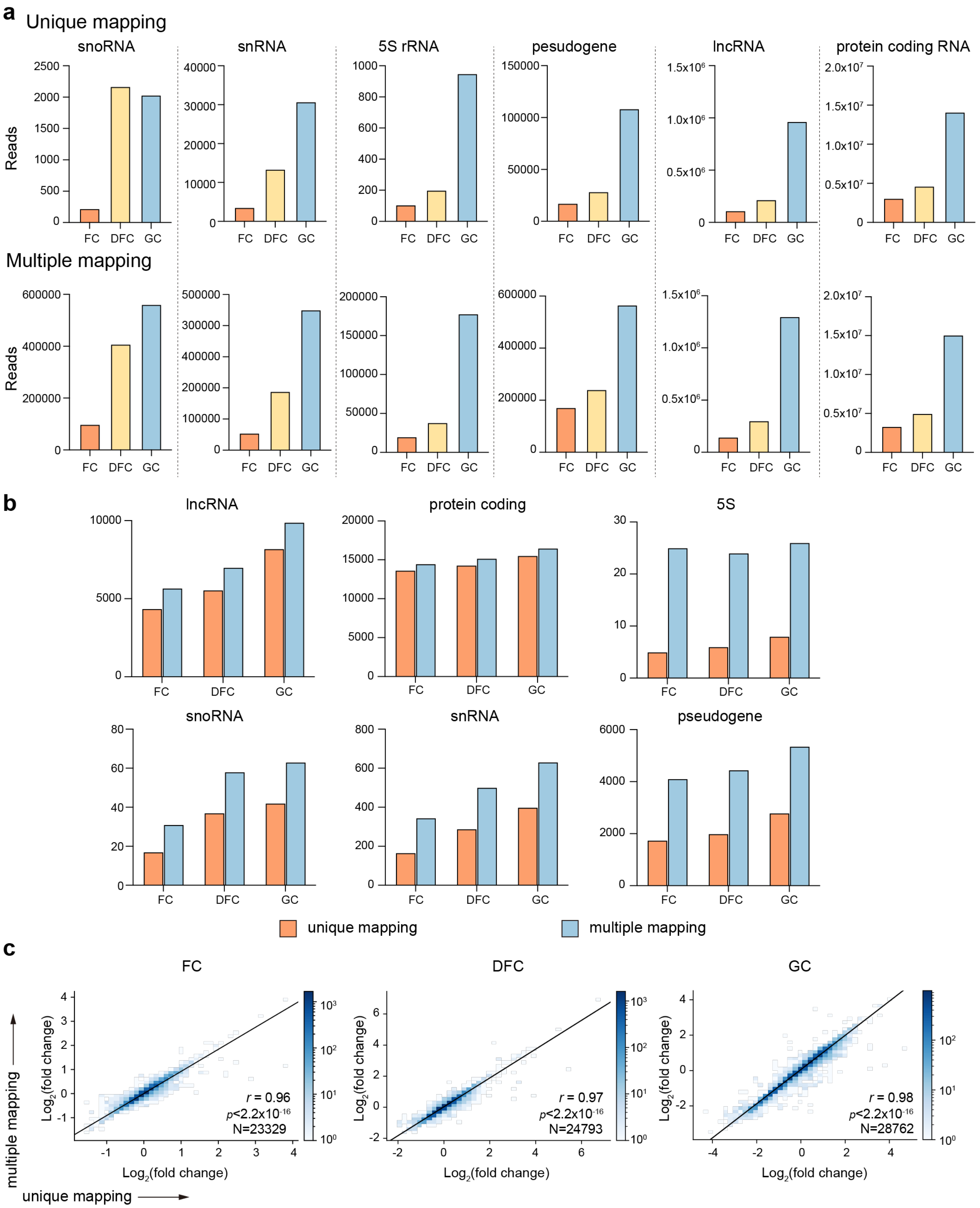


**Supplementary Fig. 2 | Comparison of unique and multiple mapping to the genome. a**, Comparison of the numbers of reads mapped to six RNA types between unique mapping and multiple mapping methods. **b**, Comparison of the number of identified genes with two mapping methods. **c**, Comparison of $Log_2$(fold change) values from DESeq2 analysis using two mapping methods. N reports the number of genes. *r* reports Pearson's correlation coefficient.

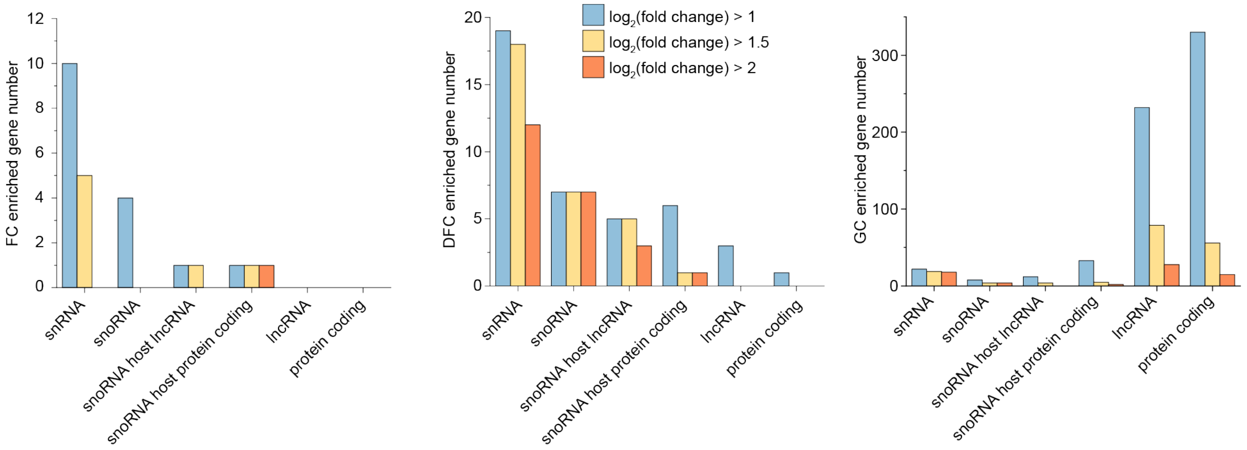


**Supplementary Fig. 3 | Numbers of genes enriched in three subdomains with increasing enrichment thresholds for different transcript types.** The criteria of identified genes were selected as adjusted $p$-value < 0.05, lfcSE < 1 and increasing $\log_2$(fold change) from 1 to 2.
